# Antarctic geothermal soils exhibit an absence of regional habitat generalist microorganisms

**DOI:** 10.1101/2024.06.06.597824

**Authors:** Stephen E. Noell, Jaber Abbaszadeh, Huw Richards, Marie Labat Saint Vincent, Charles K. Lee, Craig W. Herbold, Matthew B. Stott, S. Craig Cary, Ian R. McDonald

## Abstract

Active geothermal systems are relatively rare in Antarctica and represent metaphorical islands ideal to study microbial dispersal. In this study, we tested the macroecological concept that high dispersal rates result in communities being dominated by either habitat generalists or specialists by investigating the microbial communities on four geographically separated geothermal sites on three Antarctic volcanoes (Mts. Erebus, Melbourne and Rittman). We found that the microbial communities at higher temperature (max 65□) sites (Tramway Ridge on Erebus and Rittmann) were unique from each other and were dominated by a variety of novel *Archaea* from class *Nitrososphaeria*, while lower temperature (max 50□) sites (Western Crater on Erebus and Melbourne) had characteristically mesophilic communities (*Planctomycetes, Acidobacteriota,* etc) that were highly similar. We found that 97% of the detected microbial taxa were regional habitat specialists, with no generalists, with community assembly driven by high dispersal rates and drift (25 and 30% of community assembly, respectively), not environmental selection. Our results indicate that for microbial communities experiencing high dispersal rates between isolated communities, habitat specialists may tend to out-compete habitat generalists.

## Introduction

In ecology, the niche concept is commonly used to group organisms based on the range of different habitats they live in. Habitat generalists (which tend to be metabolic generalists (Barberán *et al*., 2014; Bell and Bell, 2021; Chen *et al*., 2021a; Malard and Guisan, 2023)) are adapted to thrive in a broader range of different ecosystems compared to habitat specialists, although some organisms may be specialized to a habitat in one area but have a broader distribution beyond the scope of the current study (Malard and Guisan, 2023). Habitat generalists that have a global distribution are usually called “cosmopolitan” to distinguish them from other habitat generalists (Herbold, Lee, *et al*., 2014). In most environments, the diversity of habitat specialists is greater than generalists, but habitat generalists tend to be the most abundant organisms; this is true both for microorganisms (Barberán *et al*., 2014; Zhang *et al*., 2018; Chen *et al*., 2021b; Gong *et al*., 2022) and macroorganisms (Brown, 1984; Magurran and Henderson, 2003). Exceptions to this trend are few, but in macroecology, high rates of dispersal between communities can lead to habitat specialists outcompeting habitat generalists, although the opposite has also been observed (Brown and Pavlovic, 1992). In contrast, at least two reported exceptions to this trend (habitat generalists being the most abundant) in microbial ecology have been found under situations of severe environmental selection and/or dispersal limitation: strong salinity gradients (Logares *et al*., 2013) and the mammalian gut (Green *et al*., 2016). Other environmental factors are well known to be highly important in structuring microbial communities (e.g., pH (Fierer and Jackson, 2006; Rousk *et al*., 2010), salinity (Jiang *et al*., 2022), water availability (Niederberger *et al*., 2019)), raising the possibility that environmental factors may be the most important factor determining the proportion of generalists and specialists in environments. In microbial ecology, it is unknown how high dispersal rates would influence the direction of community dynamics: would it result in habitat specialists finding their optimal niches and outcompeting all generalists, resulting in no generalists? Or, would it lead to an enrichment in habitat generalists that are capable of filling any niche into which they are dispersed, given their strong ability to survive dispersal (Sriswasdi *et al*., 2017; Schiro *et al*., 2022)? It is challenging to answer this question, however, given the complexity of most systems (many routes of dispersal, ecological disturbances, the influence of macroorganisms, etc.).

Antarctic geothermal regions offer a unique opportunity to resolve the outcome of high dispersal rates on the proportions of habitat generalists and specialists in a microbial community for several reasons. First, Antarctica is one of the windiest places on earth (Parish, 1988), which is likely to result in strong dispersal of microorganisms between sites within Antarctica, as has been found in the McMurdo Dry Valleys (Archer *et al*., 2019) and other areas of Antarctica (Hughes *et al*., 2004; Pearce *et al*., 2010). Second, aerial dispersal is the only possible route of dispersal between Antarctic geothermal regions, given the absence of aquifers and bird traffic connecting regions, which reduces complexity. Third, Antarctic geothermal regions have relatively simple communities, with little to no macroscopic life present (Herbold, Mcdonald, *et al*., 2014). Finally, Antarctic geothermal regions serve as refugia for thermophiles, given the vast amount of ice and snow separating regions and the extremely cold air conditions in Antarctica, resulting in little regional influence of non-thermophiles on the microbial communities present. Interestingly, past studies of thermophiles, mainly in hot spring environments, have found high niche specialization for temperature or pH, with environmental selection being the primary factor influencing community assembly and low dispersal rates between sites (Weltzer and Miller, 2013; Power *et al*., 2018a, 2023; Mahato *et al*., 2019; Fernandes□Martins *et al*., 2023). It is unknown whether the strong winds in Antarctica are capable of overcoming the generally low dispersal rates for thermophiles at Antarctic geothermal regions.

The southernmost active geothermal features in the world are located in Antarctica, surrounded by vast swathes of ice and snow: Mt. Erebus, Mt. Melbourne, Mt. Rittmann, and Deception Island (Herbold, Mcdonald, *et al*., 2014). Within these geothermal areas, the interplay between cold air (as low as −60□) and hot soils results in a large variety of unique features such as ice caves, ice chimneys, and ice hummocks. Of these regions, Mt. Erebus is the highest (3794m above sea level, a.s.l.) and most volcanically active. Two of its largest geothermal sites are Tramway Ridge and Western Crater, which have differing maximal temperatures (65□ vs 50□), geothermal features (many active fumaroles in bare soil producing large amounts of steam vs. warm soil mostly covered by ice hummocks with less steam), physicochemistry (e.g., low pH vs. high pH), and productivity levels (visible phototrophic mats and moss beds vs. primarily chemotrophic driven) (Noell *et al*., 2022). Mt. Melbourne (2733m a.s.l.) is located 358 km northeast of Mt. Erebus and only has two geothermal fields, with maximal temperatures of 50□, limited fumarolic activity, and some moss beds (Broady *et al*., 1987). Geothermal activity on Mt. Rittmann (2600m a.s.l.), 450 km northeast of Mt. Erebus and 103 km north of Mt. Melbourne, consists of one steep wall with actively steaming fumaroles and hot soil (maximum 65□) that is relatively shallow due to the steep aspect (Bargagli *et al*., 1996). Given the strong maritime influence, presence of vascular plants, and off-continent location of Deception Island north of the Antarctic circle, we excluded this site from consideration in our study. For this study, we classified the two higher temperature sites with highly active fumarolic activity (Tramway Ridge on Mt. Erebus and Mt. Rittmann) as “Active” sites and the two lower temperature sites with less active fumaroles (Western Crater on Mt. Erebus and Mt. Melbourne) as “Passive” sites.

The microbial communities present at these unique geothermal regions have been incompletely characterized, especially at Mts. Rittmann and Melbourne. On Mt. Erebus, culture-independent microbial community analyses have shown that cosmopolitan, thermophilic microorganisms dominate the surface community of Tramway Ridge, while novel and endemic members of poorly understood archaeal and bacterial groups are dominant >2cm below the surface (Soo *et al*., 2009; Herbold, Lee, *et al*., 2014; Herbold *et al*., 2024). Our study comparing the communities at Tramway Ridge and Western Crater along thermal transects found distinct microbial communities, primarily driven by strong differences in pH and water content between the two sites (Noell *et al*., 2022). Studies of life on Mt. Melbourne have focused on macroscopic organisms (Broady *et al*., 1987; Logan *et al*., 2000; Skotnicki *et al*., 2001) or were culture-based (Logan *et al*., 2000; Bargagli *et al*., 2004; Pepi *et al*., 2005), with two metagenomic datasets showing a community comprised of *Proteobacteria*, *Chloroflexota*, and candidate phylum AD3 (now *Pseudomonadota*, *Chloroflexota*, and *Dormibacteraeota* respectively) (Park *et al*., 2021; Myeong *et al*., 2024). Similarly, only culture-dependent studies of the microorganisms at Mt. Rittmann have been conducted to-date (Bargagli *et al*., 1996; Logan *et al*., 2000; Allan *et al*., 2005; Poli *et al*., 2006). These high-elevation, Antarctic geothermal sites have been theorized to have acted as refugia for terrestrial Antarctic life during highly glaciated periods (Fraser *et al*., 2014).

In this study, we used a combination of 16S rRNA gene amplicon sequencing and metagenomics to resolve the proportion of habitat generalists at these sites (i.e., do we see a large group of microorganisms common to all sites?), and determine the ecological factors shaping community assembly. Given that the microbial communities at these sites are largely uncharacterized, we began by describing the taxonomy and functionality of the microbes present at these sites before answering our ecological question. We hypothesized that the lower temperature sites (Western Crater on Erebus and Melbourne) would have different microbial communities than the higher temperature sites (Tramway Ridge on Erebus and Rittmann), and that there would be a high proportion of habitat generalists across sites, driven by high rates of aerial dispersal.

## Experimental Procedures

### Site Description and Sample Collection

Soil samples were collected from sites on Mt. Erebus, Mt. Rittmann, and Mt. Melbourne, Antarctica, between 2010 and 2019. Sampling sites with coordinates are given in Figure 1. On Mt. Erebus, two sites were sampled; the first was Tramway Ridge (TR), which is the largest geothermal site on the Mt. Erebus summit and lies within Antarctic Specially Protected Area (ASPA) No. 175. TR is relatively protected from the wind, has large numbers of actively steaming fumaroles, producing gases enriched in CO_2_ and CO (Ilanko *et al*., 2019), with hot soil reaching 65□ in places, and has extensive moss and lichen mats covering 16% of the site (Management Plan For Antarctic Specially Protected Area No. 175, 2014). The second site sampled on Mt. Erebus was Western Crater (WC), which is more exposed to the wind and has very few active fumaroles and no visible moss or lichen mats; soil temperatures only reach 50□ in ice-free grounds. On the Mt. Melbourne summit, samples were collected on Cryptogram Ridge (also protected by ASPA No. 175), which, similar to WC, is comprised of ice-free hot soil spots where temperatures reach 50□. There are some patches of moss on Melbourne, and the few active fumaroles on Melbourne are covered by ice hummocks, generating CO_2_ and methane-rich gases (Skotnicki *et al*., 2001; Management Plan For Antarctic Specially Protected Area No. 175, 2014). On Mt. Rittmann, samples were collected from the single geothermally active area on the mountain, which is also within ASPA No. 175. This site is particularly steep (∼40 degree slope), with ice free hot soil reaching 63□ and many actively steaming fumaroles with gases of unknown composition, as well as a few moss patches (Management Plan For Antarctic Specially Protected Area No. 175, 2014).

**Figure 1.**
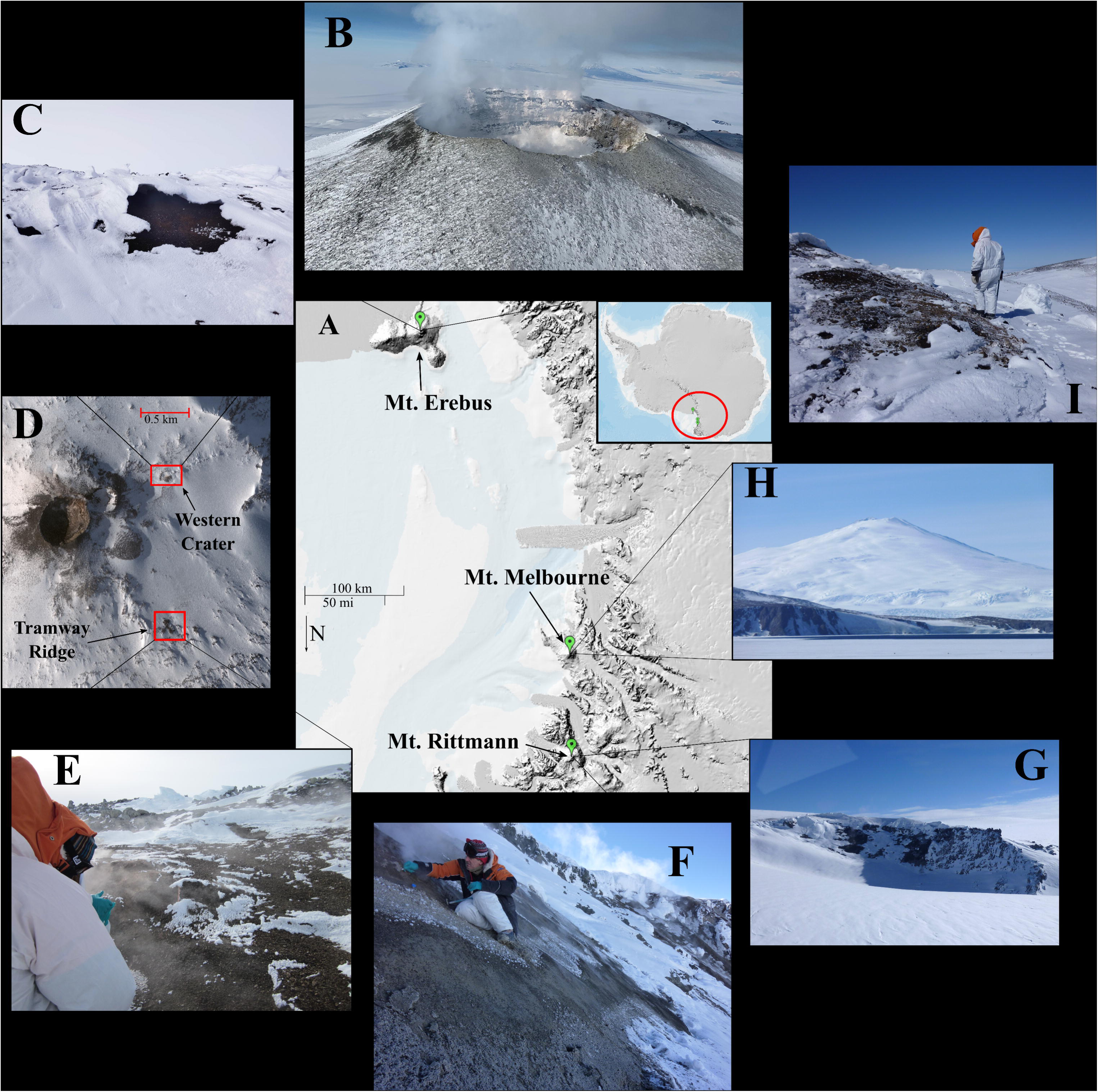
Map and images of sampling sites. (A) Satellite image of Antarctica (top right inset, red circle indicates Victoria Land); main image shows Victoria Land and Ross Island. (B) Aerial picture of Mt. Erebus main crater. (C) Picture of the hot soil site sampled at Western Crater on Mt. Erebus. (D) Satellite image showing the summit of Mt. Erebus and the two sampling sites in this study, Tramway Ridge (−77.519444, 167.116389 DD) and Western Crater (−77.519167, 167.187778 DD). (E) Image of sampling site on Tramway Ridge. (F) Image of sampling on the side of Mt. Rittmann (−73.448445, 165.497444 DD). (G) Image of Mt. Rittmann. (H) Image of Mt. Melbourne. (I) Image of sampling at a hot soil site on Mt. Melbourne (−74.350016, 164.700139 DD). Satellite image in (A) is from the Antarctic Digital Database Map Viewer https://www.add.scar.org/, Open Source, using hillshade and bathymetry layers. Satellite image in (D) was purchased from DigitalGlobe Incorporated, Longmont CO USA (2019). All other images were taken by Ian McDonald and S. Craig Cary.

Soil temperatures were measured prior to sampling using a Checktemp 1C electronic temperature sensor (Hanna Instruments, Rhode Island, United States). Mt. Erebus samples were collected along temperature transects, as detailed in (Noell *et al*., 2022). Samples were collected using a sterile spatula after first discarding the top centimetre of soil, except for one sample each at Rittmann and Melbourne where the top mat-like layer of soil was taken as the sample (Ritt3M-3 and Mel1M-2). Samples were placed into sterile 50 mL tubes and frozen for transport to the University of Waikato (Hamilton, New Zealand) for analysis. *Geochemistry*

Soil samples were analysed for geochemistry using a variety of analytical techniques. pH and electroconductivity (EC) were measured by adding MilliQ water to soil samples at a 1:2.5 soil:water ratio. pH and EC measurements were made using a Thermo Scientific Orio 4-Star Benchtop pH/Conductivity Meter (Thermo Fisher Scientific) using a 3-point calibration. For gravimetric water content (GWC) measurements, 3 g of soil sample was dried at 105□ until the sample weight remained unchanged. GWC was then calculated as the percentage of weight loss during drying (Herbold, Lee, *et al*., 2014). For elemental composition (B, Na, Mg, P, K, Ca, V, Cr, Fe, Mn, Co, Ni, Cu, Zn, Ga, As, Se, Sr., Ag, Cd, Ba, Tl, Pb, and U), dried samples (1g) were prepared and analysed as described in (Noell *et al*., 2022). For total carbon and nitrogen measurements, dried samples were also ground as for elemental analysis, then sent to Analytica (Hamilton, New Zealand) for analysis. For total organic carbon (TOC) and nitrogen (TON) measurements, samples were prepared similarly to (Noell *et al*., 2022). Briefly, 1g of samples were weighed out into acidified glass jars with cardboard disks in lids; jars and lids were washed as for the ICP-MS protocol. 1 mL 6N HCl was added to samples and dried in an oven in a fume hood for 24 hours at 95□. 100 mg aliquots of samples were analysed for TOC and TON using an automatic CN analyser.

### DNA Extraction, Amplification, and Sequencing

For 16S rRNA gene amplification as well as metagenome sequencing, genomic DNA was extracted from 0.5 to 0.9 g of each soil sample using a modified CTAB/bead beating method (Noell *et al*., 2022). To have enough DNA from low biomass samples for metagenomes, multiple replicates (up to 10) of samples were extracted and pooled. Genomic DNA was quantified using a Qubit 2.0 Fluorometer (Thermo Fisher Scientific, Massachusetts, United States).

The 16S rRNA gene (V4-V5 region) was amplified using PCR with the 515YF (5’CCATCTCATCCCTGCGTGTCTCCGACTCAGXXXXXXXXXXXXXGATGTGYCAG CMGCCGCGGTAA3’) and 926R (5’CCACTACGCCTCCGCTTTCCTCTCTATGGGCAGTCGGTGATCCGYCAATTYMTT TRAGTTT3’) primers (Caporaso *et al*., 2018; Ul-Hasan *et al*., 2019), adapted for Ion Torrent sequencing using fusion primers with a unique tag (Whiteley *et al*., 2012). PCR reaction conditions, library prep, and 16S rRNA gene sequencing are the same as described in Noell, S. E. *et al*. (2022) (Noell *et al*., 2022). 16S rRNA gene sequencing was conducted on all 21 samples on an Ion Torrent PGM (Thermo Fisher Scientific) at the Thermophile Research Unit (Hamilton, NZ).

Four samples were chosen for shotgun metagenome sequencing: Ritt1-3, Ritt3M-1 (mat sample from Rittmann), WC-50, and Mel1-3; the mat sample was included to improve the odds of recovering cyanobacterial genomes. A fifth sample (from Tramway Ridge) was previously extracted and sequenced using Illumina Hi-Seq; the two surface metagenome samples from this previous project (SRR6519253 and SRR6519256) were downloaded from the SRA using SRAtoolkit v3.0.1 and combined into one larger sequence file. For the four new metagenomic DNA samples, extracted DNA was sent to GENEWIZ Genomics Center (Suzhou, China). Next generation sequencing library preparations were constructed following the manufacturer’s protocol (NEBNext® Ultra™ DNA Library Prep Kit for Illumina®). For each sample, 1 μg genomic DNA was randomly fragmented to <500 bp by sonication (Covaris S220). The fragments were treated with End Prep Enzyme Mix for end repairing, 5’ Phosphorylation and dA-tailing in one reaction, followed by a T-A ligation to add adaptors to both ends. Size selection of Adaptor-ligated DNA was then performed using AxyPrep Mag PCR Clean-up (Axygen), and fragments of ∼410 bp (with the approximate insert size of 350 bp) were recovered. Each sample was then amplified by PCR for 8 cycles using P5 and P7 primers, with both primers carrying sequences which can anneal with flowcell to perform bridge PCR and P7 primer carrying a six-base index allowing for multiplexing. The PCR products were cleaned up using AxyPrep Mag PCR Clean-up (Axygen), validated using an Agilent 2100 Bioanalyzer (Agilent Technologies, Palo Alto, CA, USA), and quantified by Qubit2.0 Fluorometer (Invitrogen, Carlsbad, CA, USA). Then libraries with different indexes were multiplexed and loaded on an Illumina HiSeq instrument according to manufacturer’s instructions (Illumina, San Diego, CA, USA). Sequencing was carried out using a 2×150 paired-end (PE) configuration; image analysis and base calling were conducted by the HiSeq Control Software (HCS) + RTA 2.7 (Illumina) on the HiSeq instrument.

### Sequence Quality Control and Taxonomic Assignment

For 16S rRNA gene amplicon sequences, the raw reads from the Ion Torrent PGM were processed using the DADA2 pipeline with recommended Ion Torrent parameters (Callahan *et al*., 2016). Chimeras were removed using the function removeBimeraDenovo, method “consensus.” Taxonomy assignment was performed using assignSpecies with the naïve Bayesian classifier and SILVA database version 138.1. Eukaryotic, mitochondrial, and chloroplast sequences were removed and an unrooted phylogenetic tree built using the neighbour-joining method with the decipher (Wright, 2016) and phangorn (Schliep, 2011) packages in R, generating final amplicon sequence variants (ASVs).

Unfortunately, two samples (one from Tramway Ridge, TR2-64, one from Western Crater, WC-10) produced no sequence data. As these two samples had also been sequenced in a previous sequencing run (from (Noell *et al*., 2022)), we investigated whether sequence data for these two samples from the previous run could be safely used with the second sequence run samples. To control for batch effects, we tested whether other samples from the two runs had similar microbial communities and found that there was no pronounced batch effect for all samples between the two runs (Figure S1). Thus, we decided to use sequence data for TR2-64 and WC-10 from the first sequencing run alongside the second run; we denote these samples as TR2-64-1 and WC-10-1 to distinguish them from samples in the second run (TR1-32-2, WC-20-2, etc.).

For metagenome sequences, demultiplexing was performed by bcl2fastq 2.17. Raw data was filtered as follows: (1) Discard pair-end reads with adapter; (2) Discard pair-end reads when the content of N bases is more than 10% in either read; (3) Discard pair-end reads when the ration of bases of low quality (Q<20) is more than 0.5 in either read. All four metagenome samples were quality checked using FastQC v0.11.5 (Andrews) and the reports visualized using MultiQC (Ewels *et al*., 2016). Illumina adaptors were trimmed using trimmomatic v0.39 (Bolger *et al*., 2014) and sequences were trimmed for quality using the following command:

trimmomatic PE -phred33 <input files>> ILLUMINACLIP:NexteraPE-PE.fa:2:30:10:2:True MINLEN:70 SLIDINGWINDOW:4:15

In light of the large amount of duplicate sequences observed in some samples (especially the Rittmann sample) and recent evidence that suggests deduplication is beneficial (Zhang *et al*., 2022), we used fastp v0.23.2 (Chen *et al*., 2018) to deduplicate all metagenome sequences using the following command:

fastp –in1 <input file forward reads> --out1 <output forward reads> --in2 <input reverse reads> --out2 <output reverse reads> -A -D -G -Q -L –html <summary file>

Reads with ambiguous bases were removed using prinseq-lite v0.20.4 (Schmieder and Edwards, 2011) using the -ns_max_n 0 command.

### Metagenome analysis, assembly, and binning

We assessed the taxonomy of the quality trimmed metagenomes at the read level using singleM v0.13.2 (https://github.com/wwood/singlem) with the singleM pipe command; we saw no differences when we used the raw, non-quality trimmed sequences or the quality trimmed sequences (data not shown). We used Nonpareil v3.401 (Rodriguez-R and Konstantinidis, 2014; Rodriguez-R *et al*., 2018) to assess the fraction of community diversity sampled by the metagenome.

For metagenome assembled genome (MAG) assembly, binning, and processing, we used the metaWRAP pipeline v1.3.2 with default parameters (Uritskiy *et al*., 2018). We used Metaspades (Nurk *et al*., 2017) as the assembly program within metaWRAP with default settings. To minimize the amount of ambiguous bases in the assemblies, we used the non-scaffolded contigs for binning; these non-scaffolded contigs produced bins that were similar in number and quality to the scaffolded contigs (for scaffolded vs non-scaffolded, the bins produced had 94.1% vs 94.0% average completeness, 1.5% vs 1.6% average contamination, 63% vs. 64% average GC, 41309 vs. 40696 average N_50_). We used prodigal v2.6.3 in metagenome mode (Hyatt *et al*., 2010, 2012) to predict coding genes from the non-scaffolded assemblies; these predicted coding sequences were used for the metagenome functional analysis. For binning contigs in the metaWRAP pipeline, we used metaBAT2 (Kang *et al*., 2019), MaxBin2 (Wu *et al*., 2016), and CONCOCT (Alneberg *et al*., 2014). For the bin_refinement module of metaWRAP, we used targets of 80% and 5% for completeness and contamination, respectively. Bins were then re-assembled and cleaned with MAGpurify v2.1.2 (Nayfach *et al*., 2019) using the phylo-markers, clade-markers, gc-content, known-contam, and clean-bin modules to produce the final MAGs. The final statistics for these MAGs were produced using checkM v1.0.12 (Parks *et al*., 2015) with the lineage_wf command. We used GTDB-Tk v2.1.1 (Chaumeil *et al*., 2022) for taxonomy assignment of MAGs (GTDB database R214) and both Prodigal and Prokka v1.11 (Seemann, 2014) for predicting coding gene regions. We predicted optimal growth rate of MAGs using gRodon (Weissman *et al*., 2021) and optimal growth temperature using Tome v1.0.0 (Li *et al*., 2019). Finally, we used pfam_scan (https://github.com/aziele/pfam_scan) to predict pfam domains in MAGs (pfam release 35.0).

We calculated the presence/absence of MAGs across sites by utilising MinHash sketching techniques implemented with sourmash (Brown and Irber, 2016). Specifically, we created sourmash signatures of all MAGs and quality filtered metagenomic read sets from all four sites using a k-mer size of 21 and scaled to 1000. These settings allowed us to query for the presence of a MAG within a site with greater sensitivity than by using a higher k-mer or scale value. We selected these metrics to minimize the false negative of not finding a MAG that is present, though we acknowledge this possibly increases the false positive, something recently found by other groups (Koslicki *et al*., 2023). Finally, we included abundance weighting in the signature creation step of the MAG and metagenome signatures so we could measure relative abundances of MAGs present. To query presence/absence, we used sourmash *gather* on each MAG against each metagenome to calculate containment scores. These scores are a proxy for Average Nucleotide Identity where a containment score of 0.2 is equivalent to an ANI of 0.95 for a signature created with a k-mer of 21. We therefore concluded that a MAG was present within a metagenome if its score was greater than 0.2, to account for the varying completeness levels of the MAGs and to again minimize the false negative rate. We measured MAG abundances within each site using the abundance weighted percent match between the MAG and the metagenome. This correlates to the proportion of metagenomic reads that would map to the MAG, a proxy for abundance. The final set of MAGs was deduplicated for all analyses using a 95% ANI cutoff. Where duplicate MAGs were found, the MAG from the site with its highest abundance was retained.

### Analysis of 16S rRNA Gene Sequence Data

Using 16S rRNA gene sequences, microbial community structure was assessed using PcoA of center log-ratio transformed (CLR) sequence abundance on a Unifrac distance matrix using the vegdist function. Statistical differences between sites were evaluated first by testing samples did not differ significantly in their dispersion (using an analysis of multivariate homogeneity, PERMDISP), then by using a permutational analysis of variance (PERMANOVA) within the R package vegan v2.6-2 (Oksanen *et al*., 2020). Alpha diversity was also assessed within vegan via the Shannon index. Venn diagrams were plotted using the R package ggVennDiagram v1.2.0 (Gao, 2021); a custom modification to the ggVennDiagram function was used to color the sections of the Venn Diagram by the abundance of ASVs in those sections. Heatmaps were plotted using the R package pheatmap v1.0.12 (Kolde, 2019). The random forest differential abundance analysis was conducted using the R package microeco v0.13.0 with taxa level set to “all” (Liu *et al*., 2021).

To identify physicochemical parameters that correlated with microbial community structure, the mantel test function within microeco was used with default parameters. The factors that had a non-adjusted two-sided p-value < 0.05 were plotted using a distance-based redundancy analysis (dbRDA). To test for significant differences between physicochemical parameters, the cal_diff (method = anova) and plot_diff functions within the trans_env function of microeco were used. To analyse clustering of sites by physicochemical parameters, principal coordinates analysis (PcoA) of physicochemical parameters in a Euclidian distance matrix was performed using the vegdist function in the microbiome package v.1.12.0 (Lahti and Shetty, 2017).

### Metagenome Functional Analysis

We estimated the metabolic potential of each site using a method described by Greening and colleagues (Greening *et al*., 2019). In short, we used DIAMOND v. 2.1.6 (query cover > 80%) to search quality filtered assembled ORFs of the metagenomes against custom protein databases of representative metabolic marker genes designed by the authors. The metabolic marker genes include multiple pathways or energy harvesting systems, namely: sulfur cycling (AsrA, FCC, Sqr, DsrA, Sor, SoxB), succinate oxidation (SdhA), selenium cycling (YgfK), rhodopsin (RHO), reductive dehalogenation (RdhA), phototrophy (PsaA, PsbA, energy-converting microbial rhodopsin), oxidative phosphorylation (AtpA), nitrogen cycling (AmoA, HzsA, NifH, NarG, NapA, NirS, NirK, NrfA, NosZ, NxrA, NorB), NADH oxidation (NuoF), methane cycling (McrA, MmoA, PmoA), isoprene oxidation (IsoA), iron cycling (Cyc2, MtrB, OmcB), hydrogen cycling (catalytic subunit of [NiFe]-hydrogenases, the catalytic domain of [FeFe]-hydrogenases, and Fe-hydrogenases), fumarate reduction (FrdA), carbon monoxide oxidation (CoxL, CooS), carbon fixation (RbcL, AcsB, AclB, Mcr, HbsT, HbsC), arsenic cycling (ARO, ArsC), and aerobic respiration (CoxA, CcoN, CyoA, CydA). We filtered results based on different identity thresholds: PsaA (80%), PsbA and IsoA, AtpA, ARO, YgfK (70%), HbsT (75%), group 4 [NiFe]-hydrogenases, [FeFe]-hydrogenases, CoxL, AmoA, NxrA, NuoF (60%), and all others with an identity threshold of 50%. We normalized the relative abundance of present markers to RPKM.

RPKM is recommended for relative abundance comparisons within metagenomic datasets because it normalizes the data based on both sequence depth (per million reads) and sequence length (in kilobases) (Liao *et al*., 2023). We used a three-step procedure to calculate the relative abundance of each metabolic marker in the community. First, we used quality controlled reads to normalize coverage of ORFs to RPKM using CoverM v0.6.1 contig (https://github.com/wwood/CoverM) (with parameters: -p bwa-mem -min-read-aligned-percent 0.75 -min-read-percent-identity 0.95 --min-covered-fraction 0 -m rpkm). In the second step, we aligned high-quality unassembled reads to 14 universal single-copy ribosomal marker genes used in SingleM v.0.12.1 (https://wwood.github.io/singlem/) and PhyloSift (Darling *et al*., 2014) using DIAMOND (query cover > 80%, bitscore > 40) and normalized to RPKM as above. Finally, we calculated the average gene copy number of each marker in the community by dividing the read count to the gene (in RPKM) by the mean of the read counts of the 14 universal single-copy ribosomal marker genes (in RPKM). Finally, we visualized the data using ggplot2.

### Phylogenomic Tree Analysis

For the *Cyanobacteria* phylogenomic tree, we used GtoTree v1.7.00 (Lee, 2019), with our 9 cyanobacterial MAGs along with GTDB species cluster representatives from the same three Cyanobacterial families, resulting in 217 members of *Nostocaceae*, 38 for *Leptolyngbyaceae*, and 55 for *Obscuribacteraceae* (accessed on 13 October 2023). We used the *Cyanobacteria*-specific single copy gene (SCG) set available in the GtoTree package. The SCG threshold was lowered to 35% to keep the *Obscuribacteraceae* genomes. We used four members of the Terrabacteria supergroup (*Streptomyces coelicolor*, *Clostridium botulinum*, *Chloroflexus aurantiacus*, and *Meiothermus ruber*) as the outgroups to root the tree. The tree was visualized and edited using the website Interactive Tree of Life (https://itol.embl.de/upload.cgi).

### Community Assembly Analysis

To compare the abundance of cosmopolitan and site-specific microorganisms in the sites/samples, ASVs were categorized by the number of sites they were present at, with the threshold for presence at a site being greater than 0 relative abundance within either the sample or site. MAGs were classified as present or absent as described above.

To calculate niche breadth for ASVs, we first eliminated ASVs only found at one site (1878 ASVs reduced to 832 ASVs), equivalent to a 5% sample occupancy cutoff (stricter than (Sriswasdi *et al*., 2017; Xu *et al*., 2022) but not as conservative as (Hernandez *et al*., 2023)). Next, we used the R package MicroNiche v1.0.0 (Finn *et al*., 2020), with the default settings for the limit of quantification (LOQ) threshold (q = 1.6). Levin’s B_N_ was used to detect generalist/specialists across the four sample sites, while Hurlbert’s B_N_ was used to detect generalist/specialists with regards to temperature or pH. An adjusted p-value cut-off of 0.05 was used, and thresholds for generalist/specialist were determined from the null model distribution produced by the package during nice breadth calculations. Additionally, we calculated social niche breadth scores for ASVs using the script calculate_snb.py v0.2 with default parameters (von Meijenfeldt *et al*., 2023).

To measure the strength of different ecological factors on shaping community assembly in the 16S rRNA gene sequence data, we used the following commands within the microeco package with default settings: trans_nullmodel, cal_ses_betamntd, cal_rcbray, and cal_process; this approach is based on (Stegen *et al*., 2013). To generate a distance-decay plot to look at the influence of geographic distance on the phylogenetic differences between ASVs, we calculated the geographical difference between sampling sites using the distm function (distVincentyEllipsoid option) in the geosphere v1.5-18 R package (Hijmans, 2022). To calculate biological differences in the microbial communities, we chose to use the Chao index, given the differences in the sample sizes in our data set (Chao *et al*., 2004), implemented with the vegdist function in the vegan R package.

### MAG Functional Analysis

Functional annotation of MAGs was conducted using METABOLIC-G (Zhou *et al*., 2022), using the predicted proteins from each MAG as predicted by Prodigal (as above) with default parameters. Specifically, we used the full KOFAM database and set the cut-off value to assign the presence of a module to 0.75. We categorized the metabolic categories used by METABOLIC into either Pathway (sets of genes that take a metabolite from start to end-point) or Category (contains multiple distinct, not inter-connected pathways). We counted a Pathway as being present in a MAG if at least 50% of the genes in the pathway were present, and a Category as being present if there was at least one gene in that Category present.

### Statistical Analyses

All statistical analyses were conducted in R (R Core Team, 2020), with plots produced in ggplot2 v3.3.6(Wickham, 2016) and edited for aesthetics in Inkscape (https://www.inkscape.org). Data analysis was preformed using tidyverse v1.3.1 (Wickham *et al*., 2019). The ggpubr v0.6.0 package in R was used to add the results of a one-way ANOVA test (when comparing all four sites and when data was normally distributed), a one-way Krustal-Wallis test (when comparing all four sites for non-normally distributed data), or a two-sided Mann-Whitney test (when comparing active vs. passive site types) to plots (Kassambara, 2023).

## Results

### Prokaryotic Community Diversity and Composition

To characterize the microbial communities inhabiting Antarctic geothermal sites, we extracted DNA from 21 soil samples from three geothermal areas in Victoria Land, Antarctica: Mt. Rittmann, Mt. Melbourne, and two sites on Mt. Erebus, Tramway Ridge (TR) and Western Crater (WC), for a total of four sampling sites (see Figure 1 for maps and site descriptions). Two of these four sites, Melbourne and WC, have little to no fumarolic activity (i.e., fumarolic passive) and lower maximal soil temperatures (maximum 50□). The other two sites, Rittmann and TR, have many actively steaming fumaroles (i.e., fumarolic active) and higher maximal soil temperatures (maximum 65□). Thus, we categorized WC and Melbourne as “passive” sites and TR and Rittmann as “active” sites. We used 16S rRNA gene amplicon sequencing to study the microbial communities present in these samples, generating 1,871 amplicon sequence variants (ASVs) from 926,649 reads (Table S1). We also conducted metagenome sequencing on four samples, one from each site, with almost all metagenomes reaching sequencing saturation (nonpareil C value of 1 for all except TR, which was 0.93; Figure S2B). Metagenome diversity, based on nonpareil diversity indices, were 17.8, 15.6, 15.3, and 15.2 for TR, WC, Melbourne, and Rittmann, respectively, all very low compared to other habitat microbiomes (17.5 for human-associated, 19 for lake water) (Rodriguez-R *et al*., 2018). From these metagenomes, we binned 140 metagenome assembled genomes (MAGs): 42 from Melbourne, 15 from Rittmann, 23 from TR, and 60 from WC, all with completeness >80% (except for two that dropped to 77% during cleaning) and contamination <5% (Table S2).

Overall, we found that the microbial communities at WC and Melbourne (hereafter, passive sites) were more closely related to each other than to TR and Rittmann (hereafter, active sites) (Figure 2). This was most apparent from the 16S rRNA samples, where passive sites had significantly higher alpha diversity than the active sites (Figure 2A), clustered very closely together in a principal coordinates analysis (PCoA) (Figure 2B), and were more similar to each other based on abundance patterns both at the phylum (Figure S3A) and class level (Figure S3B). MAGs from passive sites also clustered together based on similarities in abundance profiles, separate from active sites (Figure S4). However, a pairwise PERMANOVA test using the 16S rRNA gene sequence data showed no significant difference between the microbial communities at any of the sites except between TR and WC (*p* = 0.05), which could be due to the small number of samples analysed at Melbourne and Rittmann. TR and Rittmann also clustered separately from each other in the PCoA plot, indicating distinct microbial communities at the two active sites (Figure 2B).

**Figure 2.**
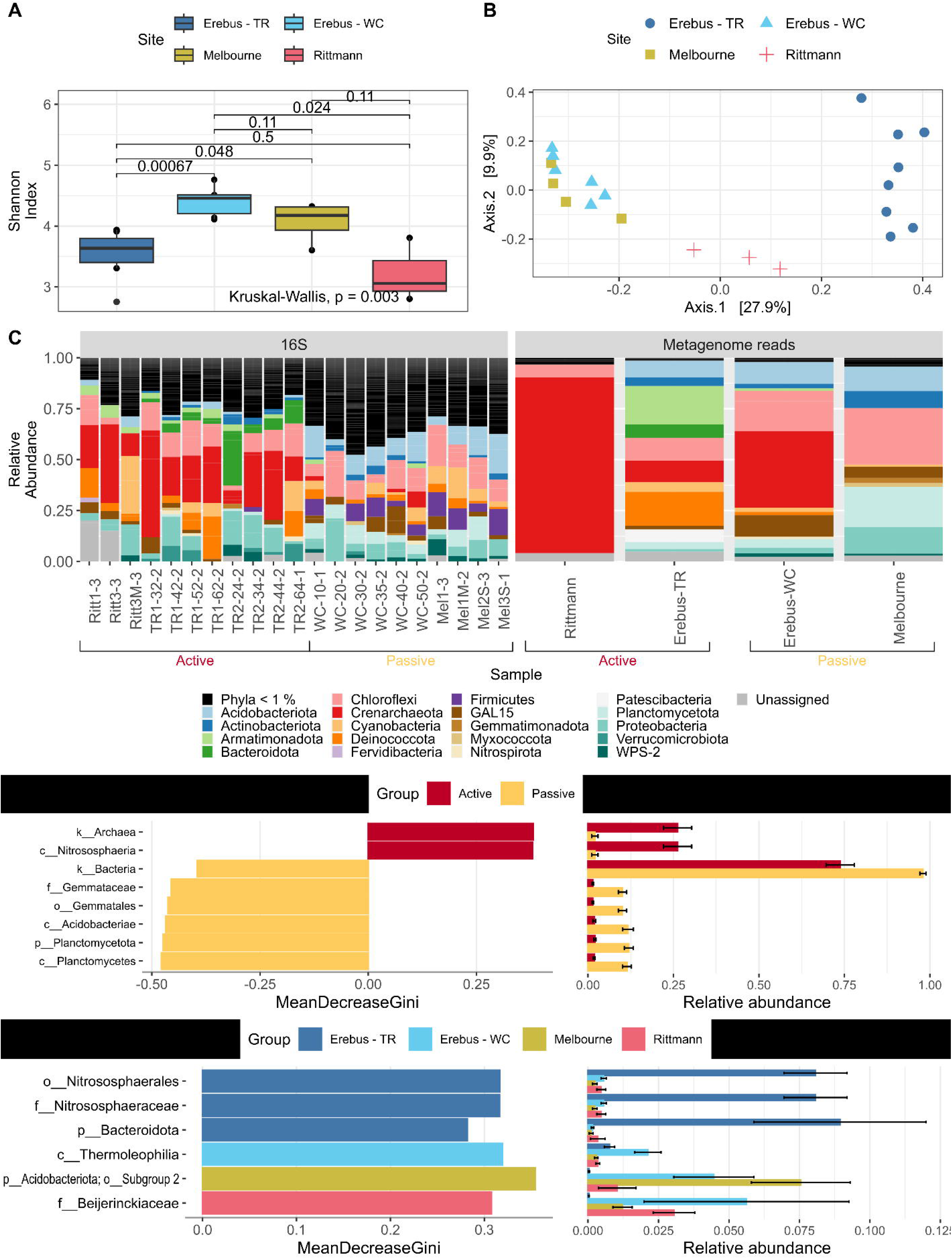
Diversity and composition of microbial communities across the four sites. (A) Alpha diversity of microbial communities based on 16S rRNA gene sequences across the four sites (Tramway Ridge (TR), Western Crater (WC), Melbourne, or Rittmann, n = 8, 6, 4, or 3 respectively) using the Shannon index of diversity. Paired significance test used an undirected Wilcox test. (B) Principal coordinates analysis (PCoA) of the microbial communities based on Unifrac similarities between 16S rRNA gene sequences. The percentages in brackets for each axis indicate the percentage of the variation in community composition explained along that axis. (C) Microbial community composition at the phylum level for both 16S rRNA gene sequencing samples (16S) and metagenome samples at the read level (Metagenome reads); singleM was used for determining taxonomy for the metagenome reads. For the 16S rRNA gene sequence data, the sequencing run used for Tramway Ridge (TR) and Western Crater (WC) samples are denoted by the last digit, either 1 for the first (data from (Noell *et al*., 2022)) or second sequencing run. For this plot, phylum names for metagenome reads were changed to match SILVA 16S rRNA gene taxonomy to allow for ease of comparison between the two methods (*Thermoproteota* to *Crenarchaeota*, *Chloroflexota* to *Chloroflexi*, CSP1-3 to GAL15, *Dormibacterota* to *Chloroflexi* (class AD3), *Eremiobacterota* to WPS-2). (D, E) Results of a differential abundance random forest analysis to identify taxa at any level that are most important for distinguishing between (D) active and passive sites, or (E) all four sites in the 16S rRNA gene sequence data. The plots on the left give the mean decrease in Gini value, which is a measure of the importance of that taxa to distinguishing the sites. The plot on the right shows the relative abundance of those taxa at the respective sites. A threshold of either 0.2 or 0.1 relative abundance was used for plotting ASVs for the active vs. passive comparison or all four sites comparison, respectively.

Based on the 16S rRNA gene sequence-based and metagenome-based community composition of the sites, we found that differences between site types were primarily attributable to greater abundances of members of *Crenarchaeota* (class *Nitrososphaeria*) at active sites, while passive sites had significantly greater abundances of phylum *Planctomycetota* (family *Gemmataceae*) and class *Acidobacteriae* within *Acidobacteriota* (Figure 2C-D, Figure S3). When looking for distinguishing taxa at each of the four sites, we found that TR was the easiest to distinguish from all other sites based on greater abundances of phylum *Bacteroidota* and the archaeal family *Nitrososphaeraceae* present (Figure 2E). Interestingly, both the Rittmann and WC metagenomes had much greater relative abundances of *Nitrososphaeria* compared to the corresponding 16S rRNA gene sequence-based community profile (85% vs. 20% for Rittmann, 25% vs. 5% for WC for metagenome vs. 16S rRNA, respectively), which could indicate primer selection against this group.

We used our complete set of non-dereplicated MAGs, categorized by the site the MAG was binned from, to examine genomic characteristics of microbes from each site. We found that MAGs binned from active sites were distinguishable by their significantly smaller average genome size (average: 3.1 Mbp for active, 4.8 Mbp for passive) and significantly higher predicted optimal growth temperature (OGT) (average: 50.8□ for active, 38.0□ for passive); MAGs from Melbourne had the largest average genome size and the lowest predicted OGT of the four sites (Figure 3A-B). We did not see significant differences in %GC, coding density, or predicted maximal growth rate between MAGs from the two site types or the four sites (Figure S5A-B).

**Figure 3.**
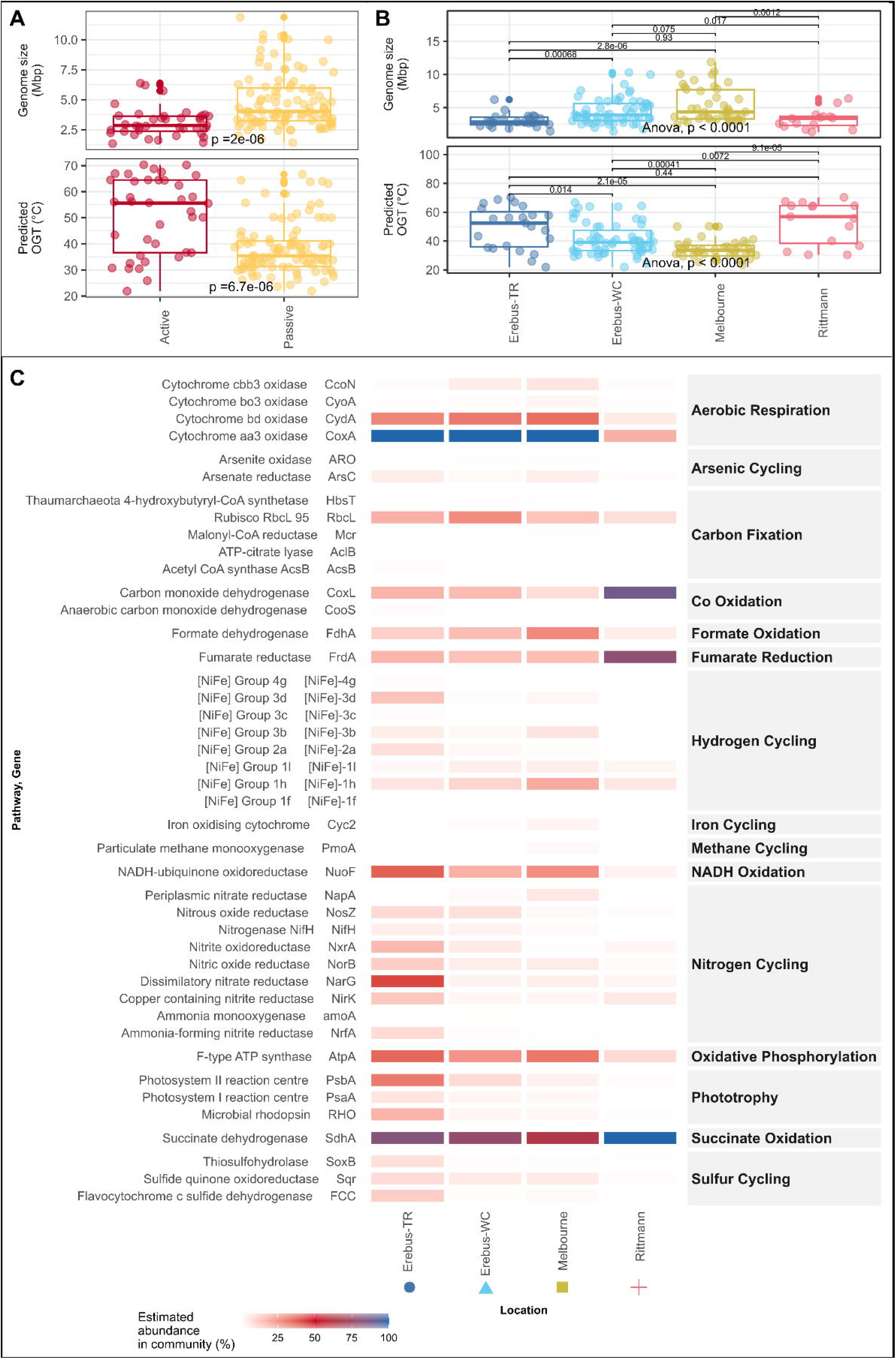
Genomic and functional characteristics of MAGs and metagenomes from each site. (A, B) Comparison of genome characteristics for MAGs recovered from different site categories (A) or sites (B). (A) n = 32 or 79 for active or passive, respectively (*p*-values: two-sided Wilcox test). (B) n = 19, 37, 42, or 13 for Tramway Ridge (TR), Western Crater (WC), Melbourne, or Rittmann, respectively. p-values given for the pairwise comparisons are for two-sided Wilcox tests. (C) Analysis of the functional potential of the microbial communities at the four site samples with metagenomes, based on the abundance of marker genes for different metabolic pathways. Abundance within community was calculated using RPKM values, which were normalized by the abundance of single copy marker genes within each metagenome. TR: Tramway Ridge. WC: Western Crater.

When we explored the physicochemical parameters that could be influencing the microbial communities at these sites, we first found that the three volcanoes clustered separately from each other in the physicochemical data (Figure S6A), with elemental composition playing a large role in this separation (Figure S6B), reflecting the unique geological origin of each volcano. When we looked for correlations between physicochemical parameters and biological communities (16S rRNA gene sequence-based), although many parameters were significantly correlated (Figure S6C), only magnesium remained significant (*p* = 0.033) after Benjamini and Hochberg adjustment. Values for selected parameters are presented in Figure S6D and all physicochemical parameter values are given in Table S3.

### Exploration of Cyanobacterial Diversity and Functions

We used the *Cyanobacteria* group as a case study to explore the influence of physicochemistry on the distribution and functioning of these important primary producers and to investigate the global uniqueness of these microorganisms. We binned an additional five cyanobacterial MAGs from the metagenome of Ritt3M-1, a cyanobacterial mat sample from Rittmann; this was necessary for our analysis as there were no cyanobacterial MAGs recovered from the other Rittmann metagenome sample. In total, we retrieved 50 cyanobacterial ASVs and 9 cyanobacterial MAGs from our data set.

Overall, we found organisms from three families (*Nostocaceae*, *Leptolyngbyaceae*, and *Obscuribacteraceae*), but could only identify a few instances of clear correlations between physicochemical parameters and the distribution and functioning of these groups (Figure 4A, B). Supporting earlier findings at the entire microbial community scale, Tramway Ridge had a distinct *Cyanobacteria* community, with species from *Leptolyngbyaceae* and the genus *Fischerella* (family *Nostocaceae*) dominant and an absence of *Obscuribacteraceae*. *Leptolyngbyaceae* was positively correlated with gravimetric water content (GWC) and negatively correlated with nickel (Figure 4C). In general, *Nostocaceae* was the most widely distributed and had a positive relationship with nickel and sulfur and was negatively correlated with GWC (Figure 4C). Only MAGs from *Nostocaceae* had the capacity to fix nitrogen (Figure S7), which matches previously measured nitrogen fixation by this group at Melbourne (Broady *et al*., 1987).We found that *Obscuribacteraceae* MAGs were non-photosynthetic, as previously observed for this group (Soo *et al*., 2014), with pathways for ethanol fermentation and fatty acid degradation instead (Figure S7B).

**Figure 4.**
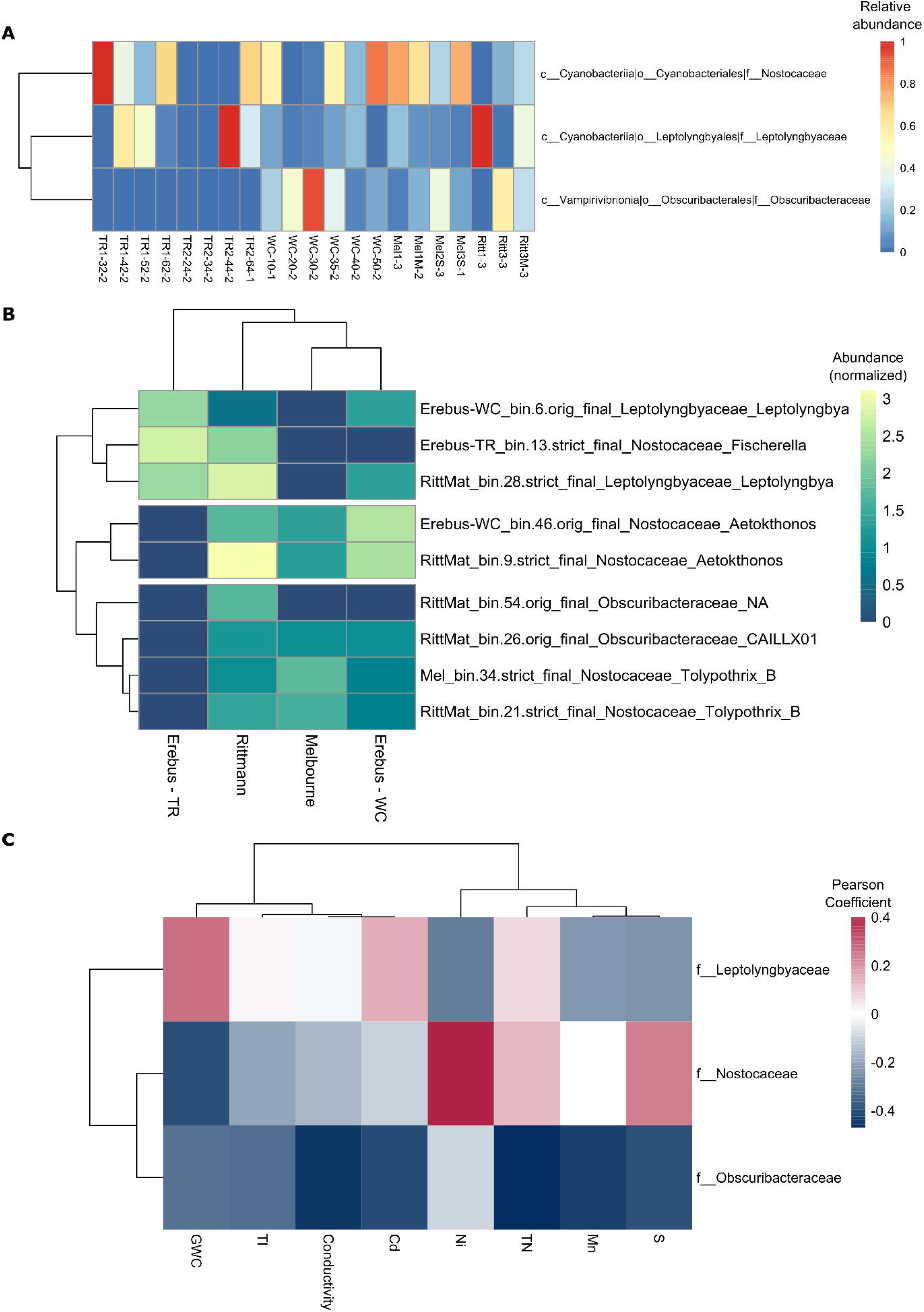
Analysis of Antarctic geothermal *Cyanobacteria* diversity and factors shaping it. Abundance heatmaps of *Cyanobacteria* communities using (A) 16S rRNA gene sequence and (B) metagenome assembled genome (MAG) data. For (A), relative abundances of *Cyanobacteria* families are presented as relative to the abundance of other Cyanobacteria. For (B), MAG abundances were marked as zero if the MAG was not present at the site. Then, the abundances were normalized: log_10_(abundance + 0.001) + 4. (C) Correlation heatmap, using the Pearson coefficient, of the cyanobacterial families to the relevant environmental variables (selected for having Mantel test p < 0.05). GWC: gravitational water content; TN: total nitrogen.

To illuminate potential evolutionary relationships between the *Cyanobacteria* MAGs from this study and globally distributed representatives of these families from GTDB (GTDB species cluster representatives, 217 *Nostocaceae*, 38 *Leptolyngbyaceae*, and 55 *Obscuribacteraceae*), we used a phylogenomic approach (Figure S8). We observed that all MAGs from our study clustered with globally distributed members of these families, from the Mojave Desert to China to the continental subsurface. The short branch lengths we observed for our MAGs also confirming the cosmopolitan distribution of these families, as has been found previously at Tramway Ridge (Herbold *et al*., 2024). These results are dependent on the completeness of the database used and should not be treated as conclusive evidence.

### Functional Comparison of Sites

We analyzed the functional potential of potentially representative microbial communities at the four sites using the community metagenome sequences, post-assembly, looking at the abundances of 52 conserved marker genes that characterize various key metabolic pathways (Figure 3C). At all sites, hydrogenases involved in atmospheric H_2_ scavenging (Group 1h and 1l NiFe hydrogenases) (Greening *et al*., 2016) were present at low abundances.

Several key differences were notable between sites. First, TR was distinguishable by an enrichment in the number of genes involved in sulfur (SoxB, Sqr, and FCC) and nitrogen (NosZ, NxrA, NirS, NirK, NorB, and NarG) metabolisms (Figure 3C). These differences are expected given the relatively large concentrations of TN and sulfur at TR (Figure S6). A notable exception to the suite of nitrogen metabolising genes detected at TR is the lack of AMO (ammonia monooxygenase) encoding genes, an exception which has recently been accounted for (Herbold *et al*., 2024). Melbourne and WC were highly similar to each other, except for the presence of methane monooxygenase genes at Melbourne and not WC (Figure 3C). Several MAGs in the genus *Methylocella* were found at Melbourne, confirming the potential for methane metabolism there, which aligns with large measured concentrations of methane at Melbourne (Geyer, 2021) (Table S2). Notably, Rittmann was distinguishable by an enrichment in marker genes for anaerobic metabolism (fumarate reductase, FrdA) and CO oxidation (CoxL), as well as much lower abundance of genes related to aerobic metabolism (cytochrome aa3 oxidase, CoxA; NADH-ubiquinone oxidoreductase, NuoF; and F-type ATP synthase, AtpA) (Figure 3C). The gas composition of Rittmann fumaroles has yet to be studied, but based on these results, is likely to be enriched in CO and depleted in oxygen.

### Habitat Specialization and Community Assembly Factors

Following our taxonomic and functional analysis of these Antarctic geothermal sites, we asked whether there was a large community of shared microorganisms between these sites (habitat generalists) and, if so, whether dispersal between sites was an important factor in structuring these communities. To do this, we first classified 16S rRNA gene sequence ASVs and MAGs by the number of sites or (for ASVs) samples at which they were present, after confirming that all samples were similarly sampled using rarefaction (Figure S2). Based on this classification, we observed that most ASVs (90%) and MAGs (77%) in our study were only found at either one or two sites/samples (Figure S9A-C, F, I). However, these site/sample-specific ASVs and MAGs comprised only a small fraction of the total read count at all samples. Instead, it was the ASVs and MAGs that were found across all four sites (36 ASVs, 3% of all ASVs; 2 MAGs, 2% of all MAGs) that were generally the most abundant (>30% of all reads for both ASVs and MAGs) (Figure S9A, B, D, G, J). Melbourne and WC shared the most ASVs and MAGs (120 ASVs, 9% of all ASVs; 36 MAGs, 32% of all MAGs) (Figure S9A, B), consistent with their similar overall microbial community composition. TR shared very few ASVs with any other site (Figure S9A, B).

Next, we asked whether these common microorganisms were adapted to thrive at all of the sites (habitat generalists), or if their presence at all sites was a product of stochastic processes. We used a robust implementation of Levin’s niche breadth (Finn *et al*., 2020) with our 16S rRNA gene sequence data (reduced to 832 ASVs by eliminating ASVs only present in one sample) to identify specialists and generalists. Levin’s niche breadth (*B_N_*) is calculated by:

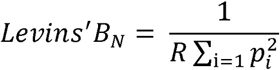

where *R* is the number of differing environments and *p_i_* is what proportion of all instances of taxon *j* is found in environment *i*. *B_N_* values close to 1 indicate broad habitat occupancy, while values close to 1/*R* indicate habitat specialization. In our data set, we defined each of our four sites as different environments and used ASV read counts for the taxon proportions.

In contrast to the theoretical distribution of niche breadth scores (Figure S10A), surprisingly, we did not find any habitat generalists, with 95% of ASVs instead identified as specialists and the rest as intermediate (Figure 5A). Because two of our sample sets (TR and WC) were collected along temperature gradients, we re-ran the analysis with only high temperature samples (>35□), but again found 96% of ASVs were specialists; only one ASV (ASV_5, GAL15) was returned as a generalist (Figure S10B). This absence of habitat generalists held true both with a relaxed limit of quantification (LOQ) for including ASVs in the niche breadth analysis, as well as when we randomly subsampled within each site to bring the sample number from all sites to the same amount (data not shown). A recently developed social niche breadth metric (SNB (von Meijenfeldt *et al*., 2023)) identifies specialists or generalists based on the compositional similarity of the communities that microbes inhabit. Using this metric, we found a distinct skew towards specialization (lower SNB score) in our data set, with a much longer tail on the specialist side both in raw SNB scores and in modified z-scores at all taxonomic ranks, supporting our findings of high habitat specialization (Figure S10).

**Figure 5.**
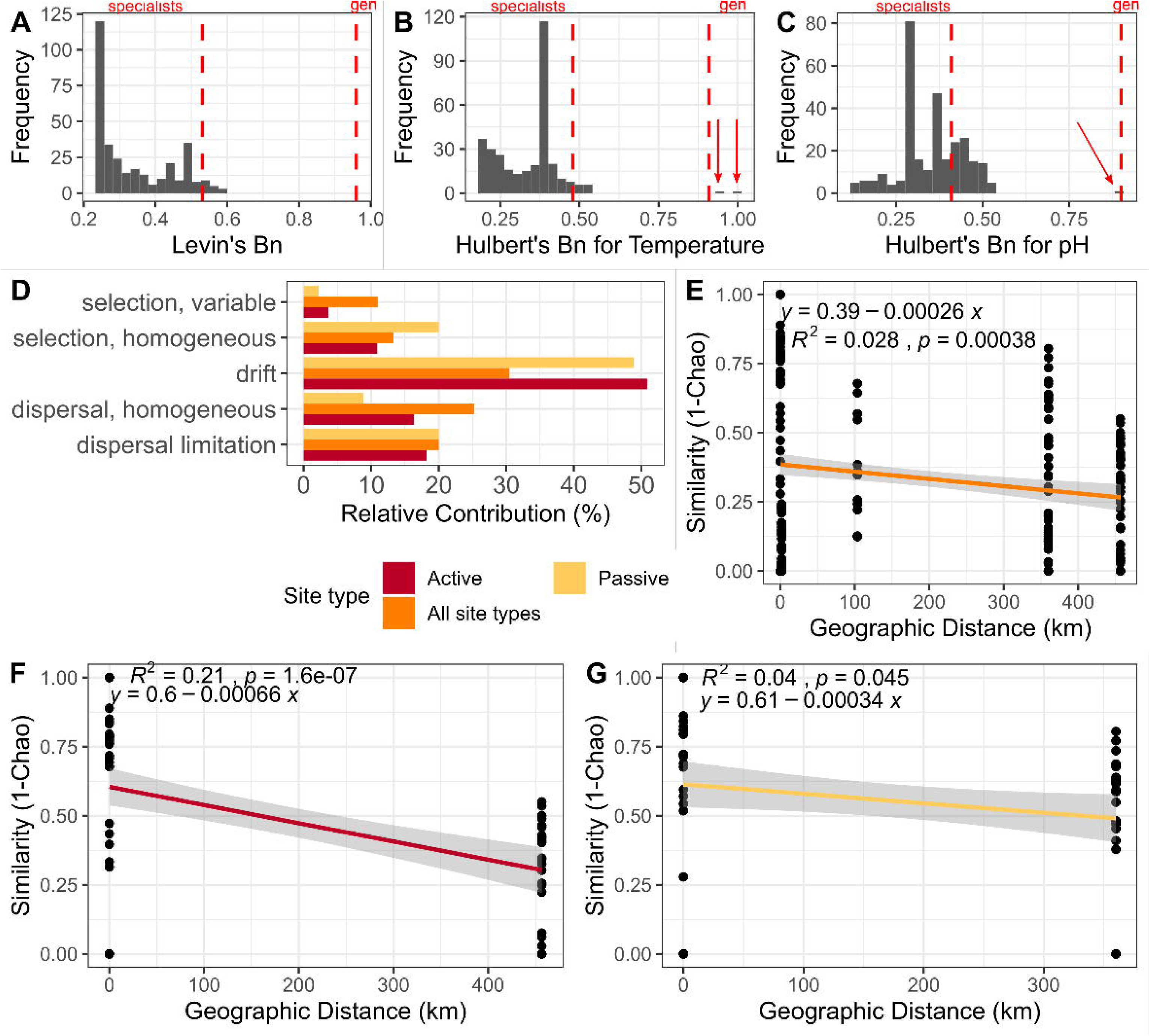
Ecological predictions for microorganisms at Antarctic geothermal sites. (A-C) Frequency plots identifying the number of either specialists or generalists for (A) habitat (using Levin’s niche breadth, B_n_), (B) temperature (using Hulbert’s niche breadth, B_n_), or (C) pH (using Hulbert’s niche breadth, B_n_) in the 16S rRNA gene amplicon sequence data. Arrows point to generalists where difficult to visualize. (D) Contribution of different ecological factors to shaping the community assembly of microorganisms in the 16S rRNA gene amplicon data. (E, F, G) Distance-decay plots for the 16S rRNA gene amplicon data with a linear regression best fit model for (E) all samples, (F) only samples from Active sites, or (G) only samples from Passive sites.

A refinement to Levins’ niche breadth adds in a measure of the resource availability across environments, called Hulbert’s niche breadth (Finn *et al*., 2020). This allows measurement of habitat specialism or generalism in reference to environmental gradients; in our case, we chose to examine temperature and pH, as these were identified as having the strongest correlations with community composition (Figure S6C). Again, we found a strong skew towards specialization for temperature and pH (95% specialists for temperature, 73% for pH) (Figure 5B, C). However, there were two ASVs identified as generalists for temperature alone (ASV 5, phylum GAL15) or both temperature and pH (ASV 42, phylum *Acidobacteriota*, genus *Bryobacter*). When examining the abundance of these ASVs across temperature and pH, they clearly had a broader distribution across different temperature/pH values than highly specialized ASVs (Figure S10E). However, their distribution within sites is more variable (Figure S10E).

Next, we explored what ecological factors might be driving the lack of habitat generalists in our data set. First, we used a common method, beta nearest taxon index and Bray-Curtis-based Raup-Crick, to calculate contribution of different factors to community assembly in the 16S rRNA gene sequence data (Stegen *et al*., 2013; Zhu *et al*., 2022; DePoy and King, 2023). We found that, across all sites, ecological drift was the strongest factor (30.5%), with homogenous dispersal (i.e., similar community structure caused by high dispersal rates) was the second strongest factor (25.5%) (Figure 5D). On the other hand, 20% of the community assembly could be explained by dispersal limitation (i.e., dissimilar community structure caused by low dispersal rates) (Figure 5D). Environmental selection (either variable selection, i.e. dissimilar community structure caused by different environmental conditions, or homogenous selection, i.e. similar community structure caused by similar environmental conditions) was the weakest factor (11.4% variable selection, 12.9% homogenous selection) (Figure 5D). Similar results were found when comparing only active sites or only passive sites, although homogeneous environmental selection was much stronger within passive sites than active (Figure 5D). The importance of ecological drift and dispersal in structuring these communities was again apparent when we examined the distance-decay plot for the 16S rRNA gene sequence data, finding a relatively flat slope of the linear regression line (*m* = −0.0003), which points to strong dispersal between sites (Figure 5E). This shallow slope held true when Active and Passive site samples were considered separately as well (*m* = −0.0007 and −0.0003 for Active and Passive, respectively) (Figure 5F, G). Slopes for similarly large-scale studies have shown much larger slopes (e.g., (Whitaker *et al*., 2003; Green *et al*., 2004; Locey *et al*., 2020)).

### Functional Imprint of Habitat Specialization

Finally, we explored the impact of high levels of habitat specialization on the microbial communities from these sites by exploring the genomic and functional profiles of MAGs based on site distribution. Generally, habitat generalists tend to be metabolic generalists (Barberán *et al*., 2014; Chen *et al*., 2021a); given the absence of habitat generalists in our data set, we predicted that the number of sites that a microbe is present at should have little to no correlation with genome size, number of unique protein domains (used as a proxy for metabolic breadth), or predicted metabolic pathways present. Rather, we would expect that most microorganisms would likely be metabolic specialists.

Indeed, we found very few significant differences (two-sided Wilcox test, Benjamini & Hochberg-adjusted p-value < 0.05) between different MAG site occupancy categories in genome size, number of unique pfam domains, or predicted metabolic pathways/categories present (Figure 6A-B). We did find that MAGs occupying three sites had significantly more metabolic pathways/categories than MAGs only occupying one site. Additionally, we observed significantly larger genome sizes and numbers of unique pfam domains in MAGs present at both WC and Melbourne compared with those occupying only WC (Figure 6B). Correlations between other genome characteristics and site occupancy were non-significant (Figure S11A-B). When we explored the ASVs that were identified previously as generalists for temperature and pH, ASVs 5 and 42, we were able to match ASV 5 to the MAG Ritt_bin.6.strict_final with 100% identity. This MAG had some hallmarks of generalism, with 20.9 predicted metabolic pathways/categories but a genome size of only 3.4 Mbp. Two MAGs (Erebus-WC_bin.32.strict_final and Mel_bin.5.orig_final) were classified to the *Bryobacteraceae* family, as was ASV 42; neither MAG contained any 16S sequence, so we cannot be sure if they match ASV 42. Still, both MAGs have genome sizes and numbers of predicted metabolic pathways/categories on the upper end of the distribution (6.2 Mbp and 10.1 Mbp, respectively for genome size; 18.2 and 26.2, respectively, for metabolic pathways/categories).

**Figure 6.**
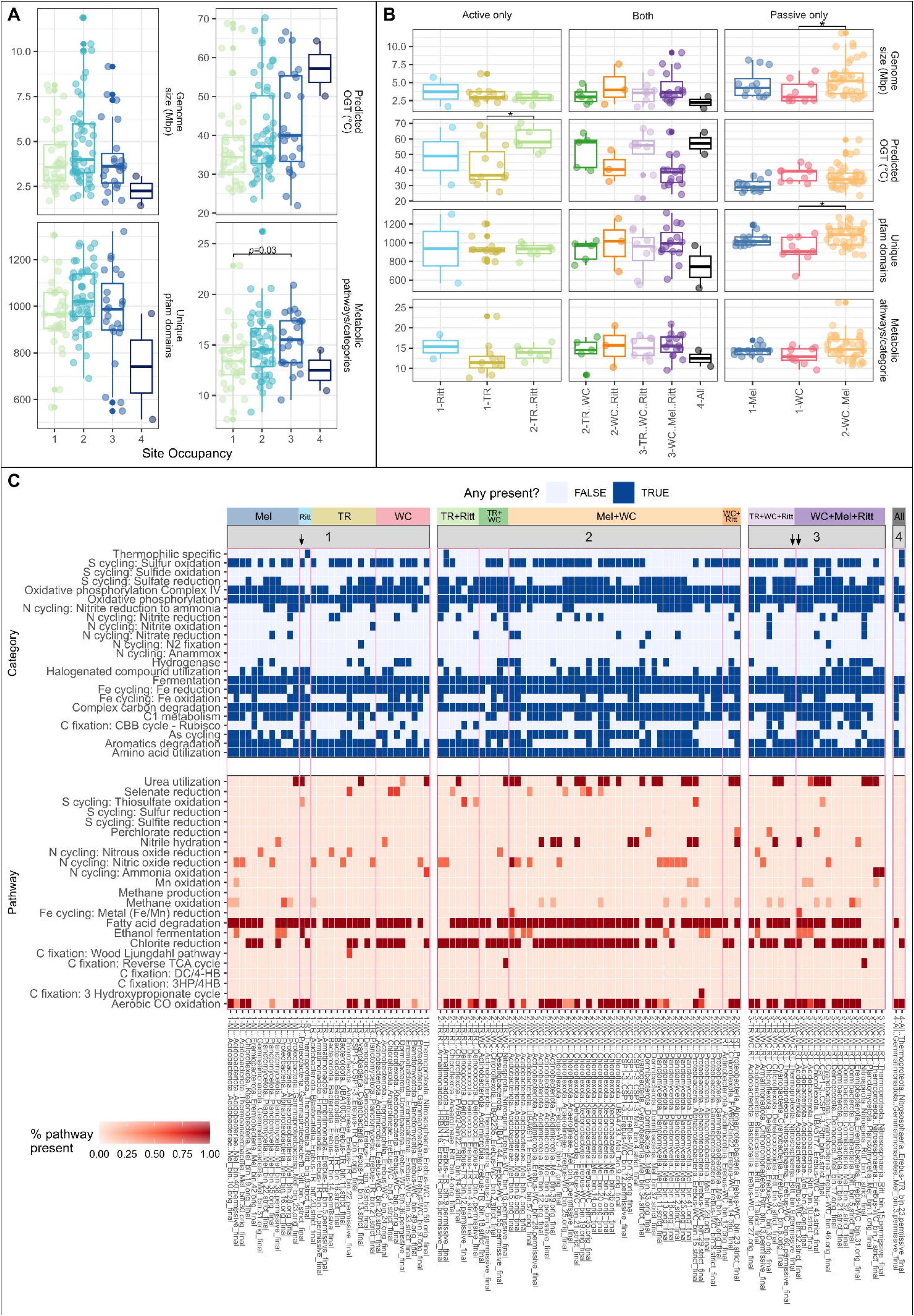
(A-B) Exploring evidence for genomic markers of metabolic generalism in metagenome assembled genomes (MAGs) from the four sites. For (A), n = 35, 52, 27, or 2 for the categories 1 site, 2 sites, 3 sites, or 4 sites, respectively. p-values for ANOVA tests conducted in (A) are, from top left clockwise order: 0.08, 0.07, 0.04, and 0.02. p-value shown on plot is the result of an undirect Wilcox test, Benjamini-Hochburg corrected. No differences for pfam domains remained significant after BH correction. For (B) and (C), n = 11, 2, or 7 for Tramway Ridge (TR) only, Rittmann only, or TR/Rittmann only MAGs, respectively; n = 5, 3, 8, 15, or 2 for TR/Western Crater (WC) only, WC/Rittmann only, TR/WC/Rittmann only, WC/Melbourne/Rittmann only, or MAGs at all four sites, respectively; n = 10, 11, or 36 for WC only, Melbourne only, or WC/Melbourne only MAGs, respectively. For (B), * represents a statistically significant (p < 0.05) result of an undirected Wilcox test after Benjamini-Hochburg correction. See Methods for how Metabolic Pathways/Categories were marked as present or absent in MAGs. (C) Heatmap showing functional predictions for MAGs based on METABOLIC predictions, separated by site occupancy (number and identity of sites occupied) and whether the metabolic predictions are categories (containing multiple, non-sequential metabolic reactions) or pathways. MAGs that are specifically mentioned in the text are indicated by an arrow. Pink boxes differentiate different site occupancy patterns. X-axis MAGs are clustered first by site occupancy pattern, then alphabetically by Phylum-Class. TR: Tramway Ridge. WC: Western Crater. RT: Rittmann. ML: Melbourne.

We investigated the different kind of metabolisms present in MAGs based on site occupancy patterns and found few patterns (Figure 6C, Table S4). We did observe a trend towards MAGs present at two or three sites being more likely to have the capacity for nitrile hydration (6%, 17%, or 11% of MAGs present at one, two, or three sites, respectively) (Figure 6C). Nitrile hydration provides microorganisms with another avenue of acquiring carbon and nitrogen needed for growth (Fang *et al*., 2015). Notably, the two MAGs that were present at all four sites lack many pathways/categories that were found in other MAGs, such as ethanol fermentation, nitric and nitrous oxide reduction, urea utilization, hydrogenases, and halogenated compound utilization, among others (Figure 6C), indicating that presence at all four sites does not require metabolic generalism.

As we initially hypothesized, there was no observable trend towards metabolic specialization, rather, relatively even numbers of metabolic specialists and generalists across sites and site occupancy patterns (Figure 6C). Rather, there were relatively even numbers of metabolic generalists (many types of pathways, aerobic/anaerobic, multiple energy sources, etc.) and metabolic specialists between the different site occupancy patterns. One example of a metabolic generalist, the MAG with the second largest number of metabolic pathways present, Ritt_bin.7.strict_final (*Gammaproteobacteria*, genus *Paraburkholderia*), appears to be capable of, among other things, urea utilization, thiosulfate and sulfur oxidation, nitrite reduction to ammonia, fatty acid degradation, ethanol fermentation, some types of complex carbon degradation and C1 metabolism, carbon fixation (via RuBisCo), and aerobic CO oxidation, yet was only present at Rittmann (Figure 6C). On the other hand, we observed multiple metabolic generalist MAGs that also had broad habitat ranges, such as Erebus-WC_bin.32.strict_final (*Acidobacteriota*, order *Bryobacterales*) which was found at all sites except TR and is capable of both organic and inorganic metabolisms, and both aerobic and anaerobic metabolism (among other things, sulfur oxidation, nitrile hydration, nitrite reduction to ammonia, iron oxidation, fatty acid degradation, ethanol fermentation, some level of complex carbon degradation, chlorite reduction, and aerobic CO oxidation) (Figure 6C).

One factor that seems to be largely negating the trends towards metabolic generalism in MAGs with higher site occupancy is the presence in our dataset of a large number of archaeal MAGs with broad habitat ranges (present at 3+ sites) with few predicted metabolic pathways and very small genomes (mean archaeal genome size: 2 Mbp). For example, Erebus-TR_bin.23.permissive_final (class *Nitrososphaeria*), present at all four sites, has a limited metabolic repertoire (genes for aerobic CO oxidation, chlorite reduction, nitrite reduction, and sulfur oxidation), with mostly aerobic pathways (Figure 6C). Similarly, Ritt_bin.9.permissive_final (class *Nitrososphaeria*, order *Conexivisphaerales*), present at all sites except Melbourne, is capable of the same set of metabolic pathways minus nitrite reduction (Figure 6C).

## Discussion

In this research, we have characterized the microbial communities inhabiting remote, isolated Antarctic geothermal sites and investigated the impacts of high dispersal rates on the abundance and distribution of habitat specialists and generalists in a patchy environment. We found that lower temperature, passive sites (WC, and Melbourne) share very similar microbial communities that were dominated by *Plancomycetota* and *Acidobacteriota*, while the hotter, active sites (TR, and Rittmann), each harbored distinct communities with high abundances of *Nitrososphaeria*. We also discovered an absence of habitat generalists in our data set, with no microbes thriving at all four sites, driven by high dispersal between sites and high ecological drift. It should be noted that “habitat generalists” here refers to generalists across Antarctic geothermal sites, which is different from “globally cosmopolitan” microbes. We also note that only a single sample from each site was sequenced for metagenomics, which may not be sufficient to capture the community heterogeneity present at these sites.

### Site Characterizations

Both taxonomically (Figure 2, Figure S3, Figure S4, Figure S9A, B) and functionally (Figure 3), the microbial communities at active and passive sites are distinct from each other. The higher temperatures at active sites appear to be a major barrier preventing microorganisms from passive sites colonizing active sites. This is apparent from the significantly lower predicted optimal growth temperatures (OGT) of MAGs from passive vs. active sites as well as that of MAGs from the Rittmann mat sample (32□) vs. the Rittman surface sample (60□), which was collected directly underneath the Rittmann mat sample. It is likely that the steep temperature gradient between the air (generally below −30□) and hot soil (60-65□) is a large factor in facilitating microorganisms with lower temperature optima, such as *Cyanobacteria* and *Bacteroidota*, to survive at hot spots at Rittmann and TR.^103,104^ MAGs present at both active sites and at least one passive site tended to have higher predicted OGTs than all other MAGs (Figure 6B), which has been observed before for thermophiles (i.e., there is a positive correlation between maximal growth temperature and temperature growth range (Miller, Williams, *et al*., 2009; Weltzer and Miller, 2013)).

TR and Rittmann have additional, unique physicochemical properties (nutrient availability and soil depth) that appear to influence the microbial community composition present. The extensive phototrophic mats and moss beds present at TR that are not found at the other sites(Skotnicki *et al*., 2001) likely provide the complex carbohydrates required for microorganisms from *Bacteroidota* to thrive there, despite their low predicted OGT (35□). The high levels of total nitrogen (TN) at TR (likely, in part, due to the extensive moss beds observed) could explain the presence of *Leptolyngbyaceae* there and its absence from other sites, given the inability of MAGs from this family in our data set to fix nitrogen. Finally, Rittmann has lower richness than other sites, both in terms of 16S rRNA gene sequence data (Figure 2A) and number of MAGs binned (Rittmann was lowest at 15), perhaps due to the shallow layer of topsoil present caused by the steep aspect. Shallow depth to bedrock has been previously associated with lower bacterial counts and diversity (Cartwright and Hui, 2015; Cartwright *et al*., 2016). Rittmann has previously been found to have a lower diversity of *Cyanobacteria* than Melbourne and TR (Bargagli *et al*., 1996).

It is challenging to put the functional characteristics of Antarctic geothermal microbial communities in a global context, given the scarcity of metagenomics studies examining microbial communities inhabiting geothermally heated soils (not hot spring aqueous or sediment) in other parts of the world. However, compared to globally distributed hot springs (Shu and Huang, 2021), we found Antarctic geothermal systems to be notably low in sulphur and associated metabolisms (Figure S6D, Table S3). For instance, in Yellowstone hot springs, genes associated with sulphur metabolism (sulfur and sulfate reduction, reduced sulfur oxidation, and sulfite oxidation) were found to be among the most abundant metabolic genes (Inskeep *et al*., 2010), with similar results found in other geothermal systems globally (Poddar and Das, 2018; Power *et al*., 2018b; Prieto-Barajas *et al*., 2018; Wilkins *et al*., 2019). Erebus has long been known to be a low-sulphur volcano, but the reason behind this phenomenon have yet to be untangled (Sweeney *et al*., 2008; Sims *et al*., 2021). On the other hand, we observed large abundances of hydrogenases and carbon monoxide dehydrogenase genes at all our sites. Other geothermal systems globally have been observed to have similarly high abundances of genes related to metabolizing geothermally produced gases, especially hydrogen (Boyd *et al*., 2010; Lindsay *et al*., 2019; Shu and Huang, 2021). Interestingly, non-geothermally heated soils in Antarctica are known to be populated by a number of microbes that use trace gas (H_2_, CO, and CO_2_) metabolism for survival, forming the base of the food chain along with photosynthesis, resulting in similarly high abundances of these phenotypes as observed in our samples (Ji *et al*., 2017), but with a different origin for the gases (volcanic vs. atmospheric). In fact, some of the same, broad microbial groups conducting trace gas metabolism in cold Antarctic soils are present in our samples (*Ca.* Dormibacterota and *Ca.* Eremiobacterota).

### Evidence of Dispersal Leading to High Habitat Specialization

We had hypothesized that high rates of dispersal between Antarctic geothermal sites would result in a large number of shared microbes (habitat generalists) across all sites. Although we did, indeed, find evidence of high dispersal between sites (25% of all variation in our 16S rRNA gene sequence data can be explained by homogenous dispersal), we found support for the opposite of our hypothesis: there were no habitat generalists between sites, with the only shared microbial taxa present being clearly due to mass effects, not adaptation to multiple sites. This lack of habitat generalists cannot be explained by environmental selection, as variable environmental selection only accounted for 11% of community differences between all sites and there were very few environmental parameters significantly correlating with the overall biological community, and *Cyanobacteria* in particular (Figure 6C). Our results stand in stark contrast to past studies of thermophiles(Sharp *et al*., 2014; Power *et al*., 2018a; Louca, 2022; DePoy and King, 2023) and habitat specialists(Pandit *et al*., 2009; Xu *et al*., 2022) that have found environmental selection to be the primary factor influencing distribution, not dispersal, although one study did find a large influence of dispersal on the abundant biosphere of Tibetan hot springs (Zhang *et al*., 2018). We note that many thermophilic habitat generalists have been identified before, even along strong environmental gradients, e.g. (Weltzer and Miller, 2013; Wall *et al*., 2014; He *et al*., 2023; Sriaporn *et al*., 2023), so an absence of habitat generalists in our data set is surprising, although many thermophiles tend to be habitat specialists for temperature (Miller, Strong, *et al*., 2009; Mahato *et al*., 2019; DePoy and King, 2023).

Our findings provide key insights into how microbial communities assemble, with high dispersal rates and ecological drift leading to a lack of habitat generalists. These results are similar to past macroecological studies that have found specialist species are favored when dispersal is high and conditions are stable (i.e., low short-term habitat disturbance) (Odum, 1969; Wilson and Yoshimura, 1994; Marvier *et al*., 2004; Büchi and Vuilleumier, 2014; Koffel *et al*., 2022). High dispersal rates allow specialist species to reach optimal niches and outcompete generalist species. This does not preclude the existence of globally cosmopolitan microbes at these sites, as has been postulated previously (Herbold, Lee, *et al*., 2014), given that cosmopolitan species may be specialists for certain environments. Additionally, highly stable conditions like those experienced at these sites (Soo *et al*., 2009), partially a result of their isolation and the absence of outside disturbances, results in few opportunities for habitat generalists to take over niches vacated by specialists during a disturbance. High environmental stability has been found in macroecology to result in a selection towards K-strategists (lower growth rate, few offspring) (Odum, 1969), which may be a contributing factor in the slow predicted growth rates of MAGs in our data set (average across all four sites: 9.2 d^−1^, fastest: 1.9 d^−1^). The other factor influencing the lack of habitat generalists in our data set is the large effect of ecological drift (30% of community assembly), likely due to the small population size at these sites. In macroecology, ecological drift can override niche effects in small populations, resulting in a co-occurrence of two species even when one is a superior competitor (Orrock and Watling, 2010).

The high level of habitat specialization at these sites may also be indicative of high levels of endemism, as these two have been found to be correlated in macroecological studies (Cowling and Holmes, 1992; Ibarra and Martin, 2015). Antarctica is home to an unusually high level of endemic organisms due to its geographic isolation (Vyverman *et al*., 2010). Tramway Ridge has previously been found to harbor a number of endemic microbes (Herbold *et al*., 2024). Additionally, the most abundant organism across all sites was a highly divergent member of the class *Nitrososphaeria* that lacks genes for ammonia oxidation, despite this class being primarily known for its ammonia oxidation activities. The functional potential of this MAG has previously been inferred, with the proposed name *Candidatus* Australlarchaeum erebusii (Herbold *et al*., 2024). Despite its high abundance, this MAG (Erebus-TR_bin.23.permissive_final) had the smallest genome size (1.4 mbp) in our data set. Culturing efforts are needed to better understand the mechanisms by which this novel archaeon is capable of wide habitat range despite limited metabolic flexibility, especially given that *Archaea* are generally much more dispersal limited than *Bacteria* (Louca, 2022). Our hope is that further microbial explorations at these unique sites, with broader sampling at more areas and further metagenomic, transcriptomic, and culturing work, will help uncover the wealth of novel microorganisms present.

## Acknowledgments

The author(s) wish to acknowledge the contribution of NeSI to the results of this research. New Zealand’s national compute and analytics services and team are supported by the New Zealand eScience Infrastructure (NeSI) and funded jointly by NeSI’s collaborator institutions and through the Ministry of Business, Innovation and Employment. URL http://www.nesi.org.nz. We especially recognize the logistical support from Antarctica New Zealand for fieldwork on Mt. Erebus and support in the field from Jon Tyler. We would like to give a big thanks to Roanna Richards-Babbage for DNA extractions and preparing samples for ICP-MS, TC/TN, and TOC/TON.

## Data Availability

All R scripts used to analyze the data are available on Github at https://github.com/ThermophileResearchUnit/Antarctic_volcano_comparison_manuscript. 16S rRNA gene sequence data has been deposited at DDBJ/EMBL/GenBank under the accession KIGX00000000. The version described in this paper is the first version, KIGX01000000. Metagenome reads are available on SRA under BioProject ID PRJNA1113340.

