## Supplementary Material for "Antarctic geothermal soils exhibit an absence of regional habitat generalist microorganisms"

**Supplemental Tables**

**Supplemental Table 1** Sample abundances (read counts), Silva-based taxonomic predictions, and 16S rRNA gene sequences for all ASVs from this study.

**Supplemental Table 2** Names and statistics for metagenome assembled genomes (MAGs) used in this study. Taxonomic classification is according to GTDB. Abundance is the abundance calculated by sourmash, not a relative abundance. OGT: optimal growth temperature.

**Supplemental Table 3** Physicochemical parameters and elemental levels for all soil samples. ND: not detected. BLD: below limit of detection.

**Supplemental Table 4** Functional predictions for metagenome assembled genomes (MAGs) discussed in this study. Numbers given are for the number of hits for that marker gene in that particular MAG, based on the output of the METABOLIC program. Cells with values above 0 are highlighted in green.

**Supplemental Figures**

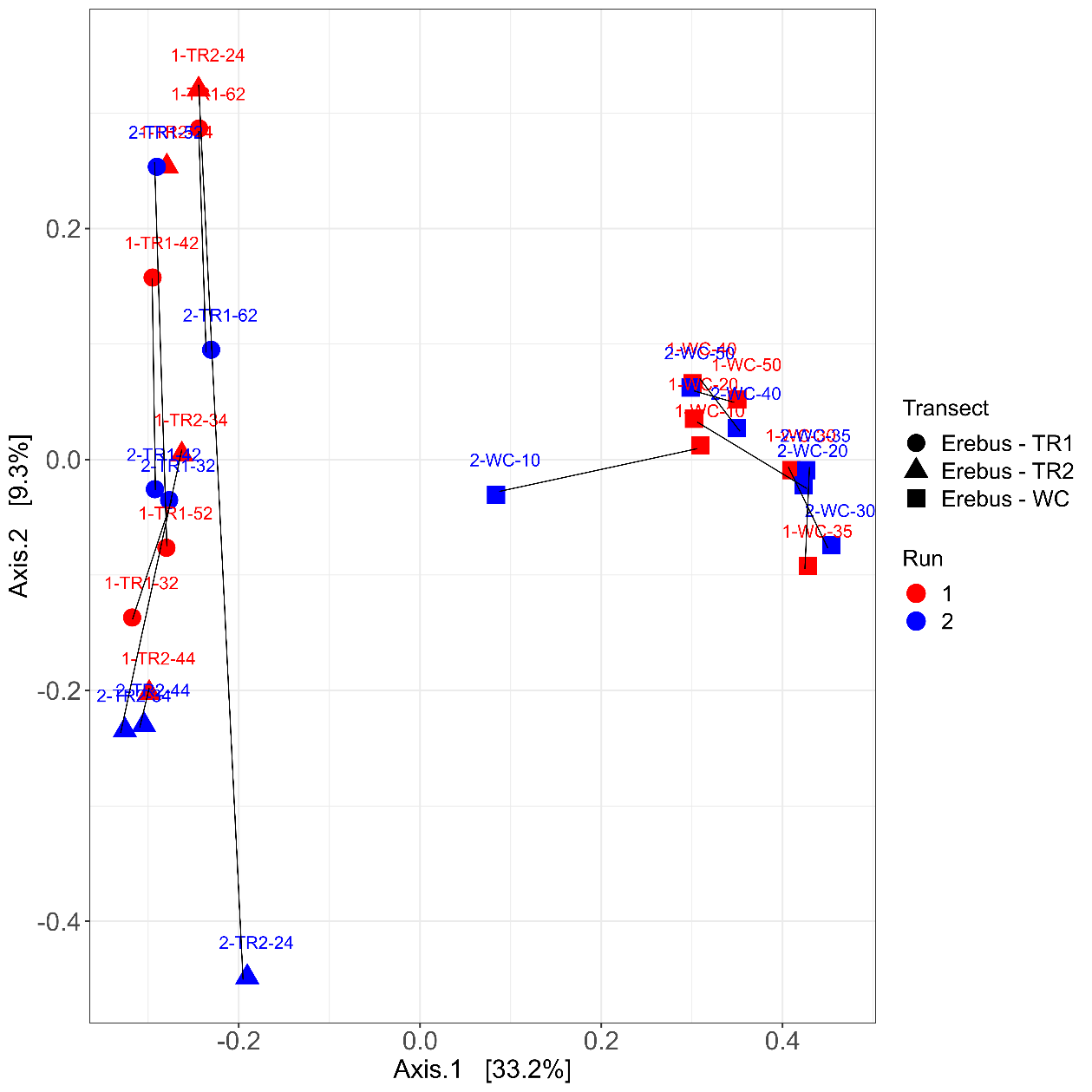

**Supplemental Figure 1** Principal Coordinates Analysis (PCoA) comparing the microbial communities (center-log ratio transformed) in all Erebus samples between the two distinct sequencing runs. Black lines show the distance between the same samples on different sequencing runs.

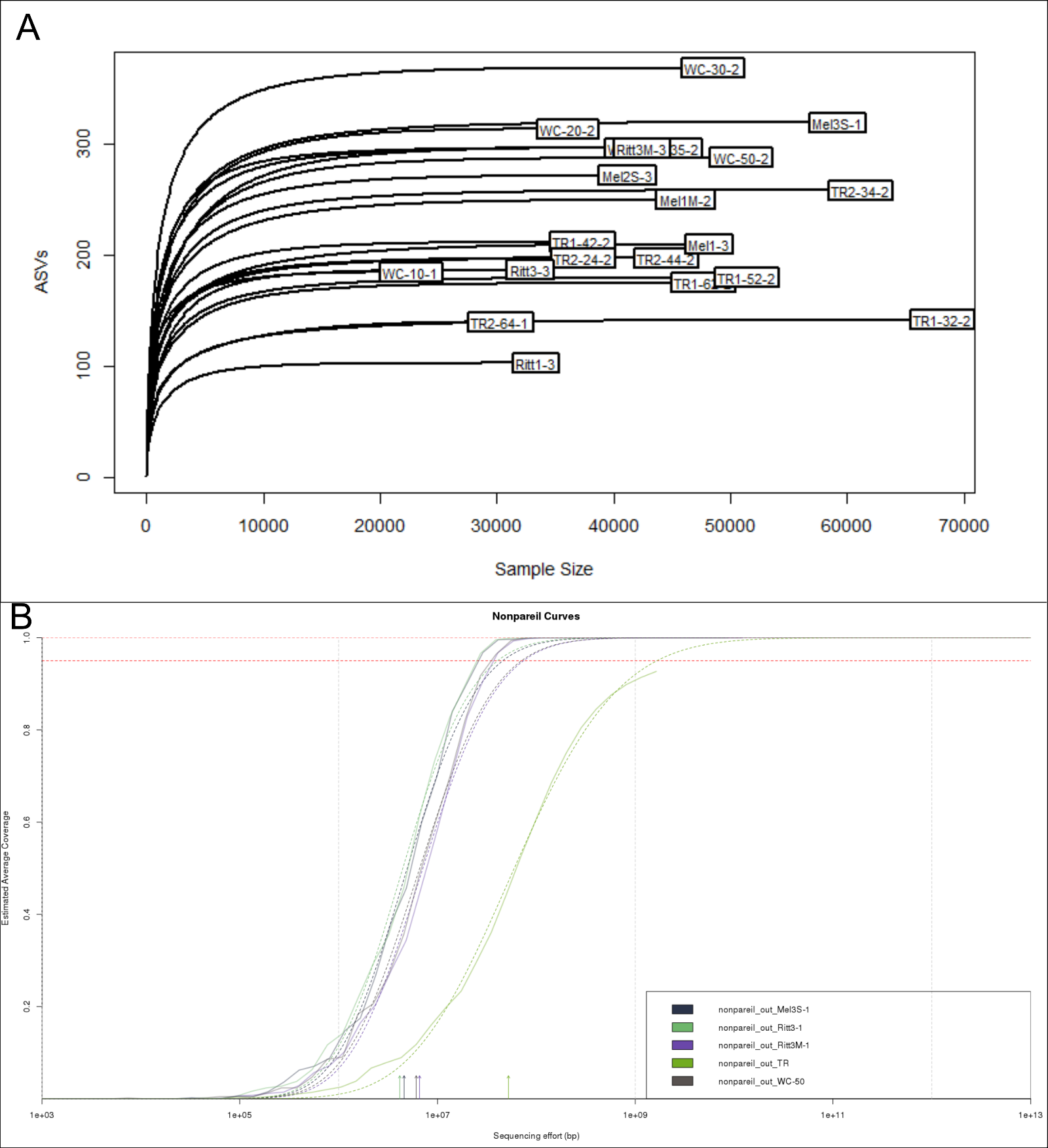

**Supplemental Figure 2** (A) Rarefaction curves for all 16S rRNA gene sequencing samples. (B) Nonpareil curves for all metagenome samples, where the solid lines indicate actual sequencing effort and the dotted lines indicate optimal sequencing effort needed to reach full sequencing depth.

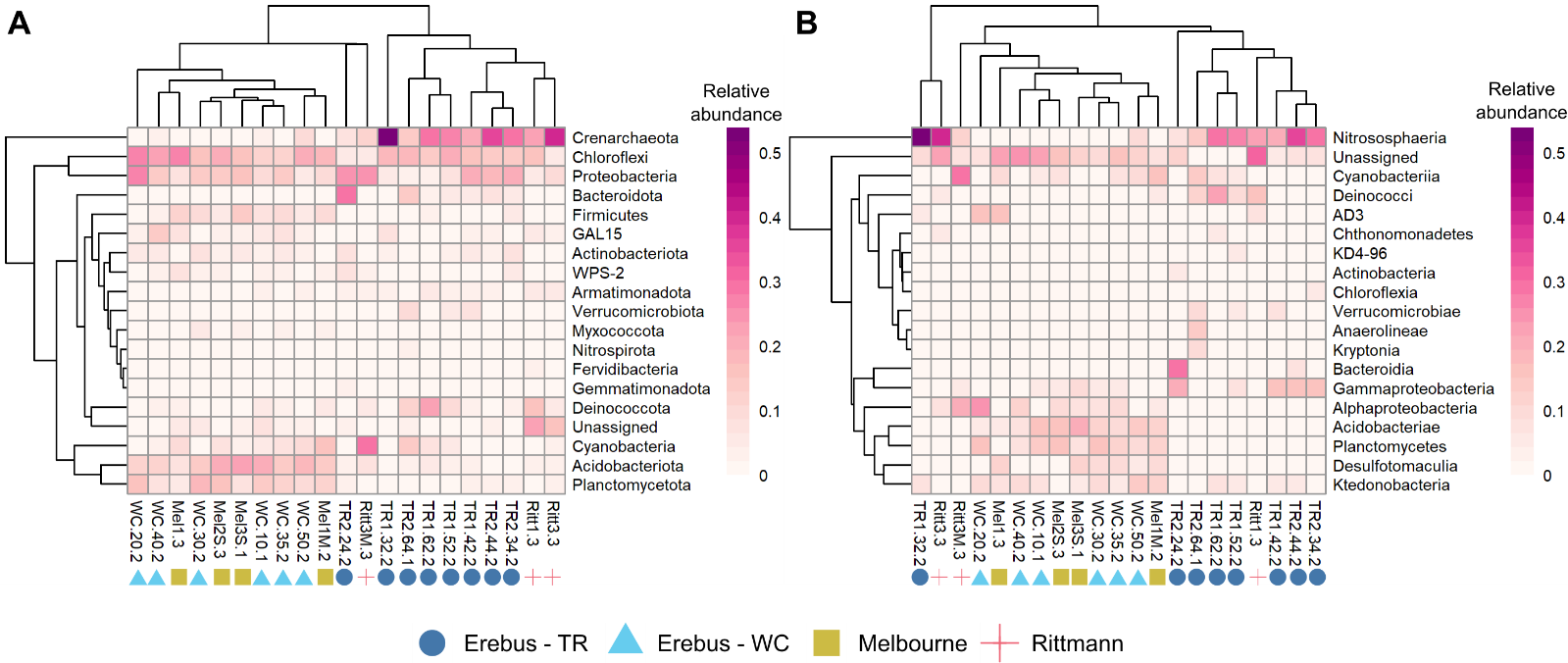

**Supplemental Figure 3** Heatmaps and dendrograms showing relative abundance of the phyla (A) or classes (B) within each 16S rRNA gene sequencing sample that had a relative abundance above 2% in at least one sample. The dendrograms do not imply an taxonomic relationship between organisms in the heatmap, only similarity in abundance patterns.

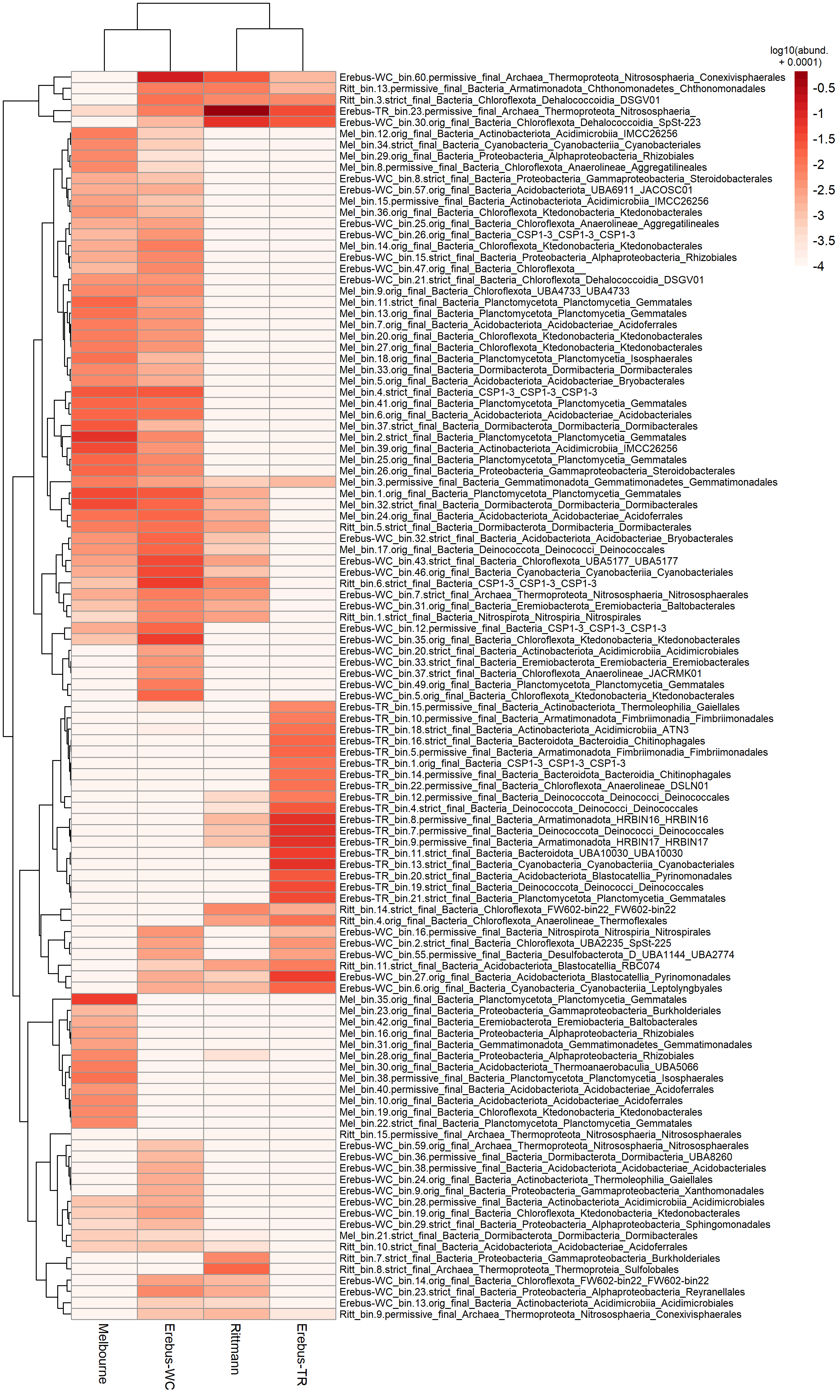

**Supplemental Figure 4** Heatmap of the abundance of metagenome assembled genomes (MAGs) from the four sites. MAGs were first defined as present/absent at a site based on sourmash containment scores from that site; the abundance of MAGs that were absent from a site were set to 0.0001 for ease of visualization and the log_10_ value of all abundances plotted. The dendograms do not imply taxonomic similarity between MAGs, only similarity in abundance profiles. The site that a MAG was binned from is given at the beginning of the MAG identifier; GTDB taxonomic information is given down to Order where available.

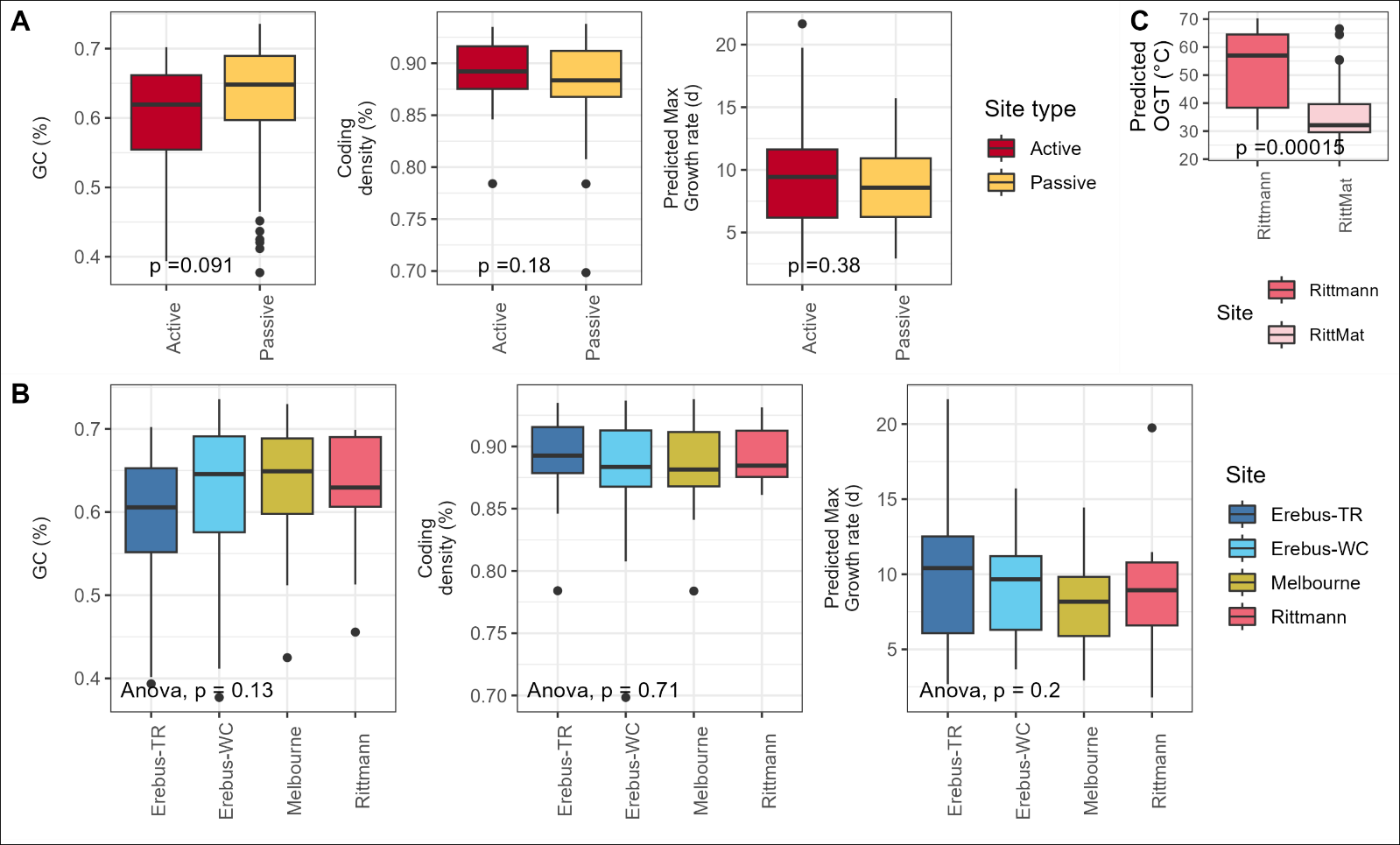

**Supplemental Figure 5** (A, B) Comparison of genome characteristics for metagenome assembled genomes (MAGs) binned from the four sites, either (A) grouped by site type (active vs. passive) or (B) all compared to each other. (C) Comparison of the predicted optimal growth temperature (OGT) of MAGs from either the Rittmann or the Rittmann mat (RittMat) sample. In (A, C), *p*-values given are the result of a two-sided Wilcoxon signed-rank test comparing group means.

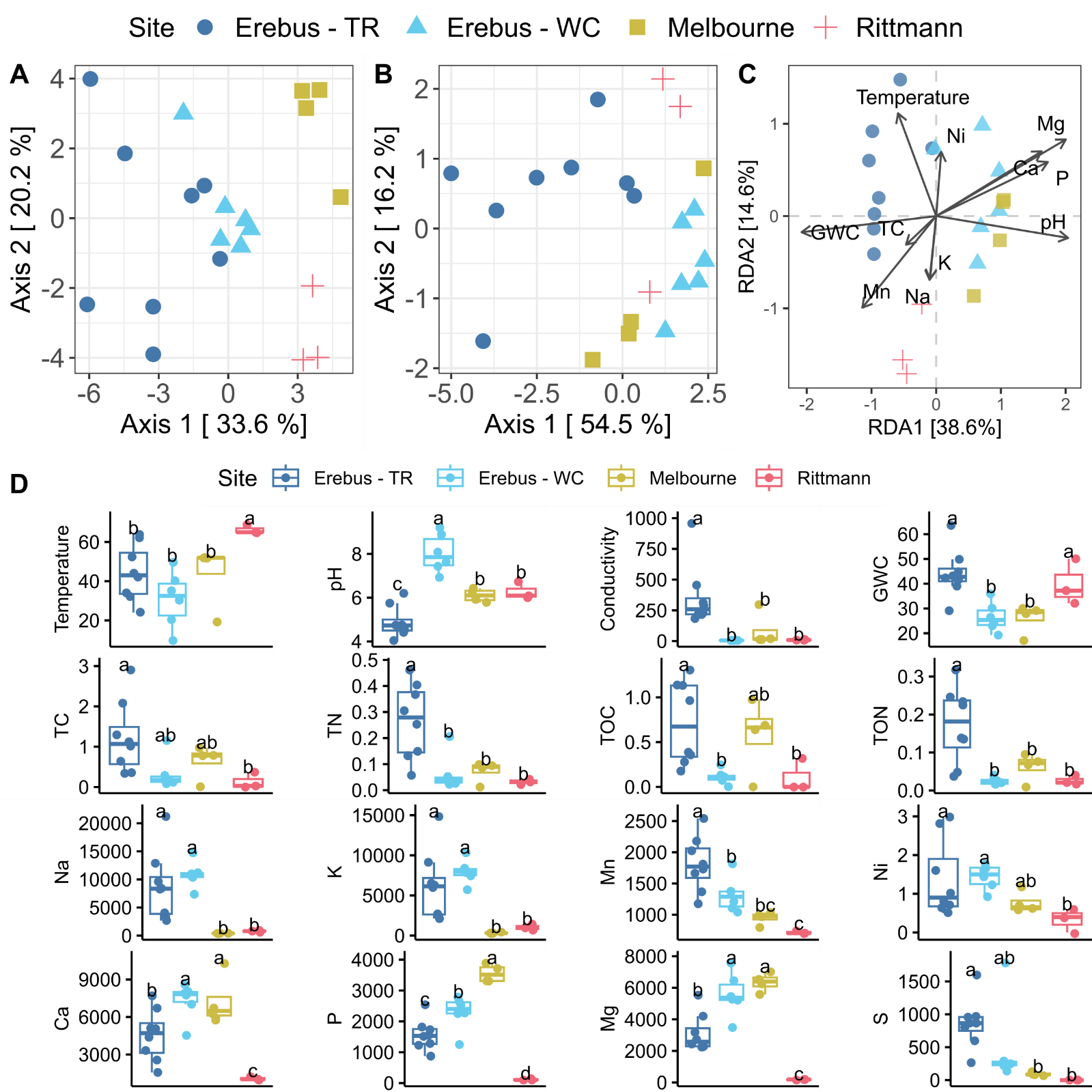

**Supplemental Figure 6** Physicochemical characterization of the four sites reveals parameters that are correlated with the microbial community present. (A) Principal Coordinates analysis (PCoA) of all physicochemical parameters (elemental and other) for each sample, using Euclidian distances. (B) PCoA of physicochemical parameters excluding elemental data. (C) Distance based redundancy analysis (dbRDA) showing the directionality of correlations between physicochemical parameters and the microbial communities present. Only parameters that were significantly correlated (Mantel test, pearson’s coefficient, p<0.05) were included. Only Mg was still significant after correcting for multiple tests (Benjamini-Hochberg correction). (D) Boxplots of basic soil physicochemical properties and elemental levels that were either significantly correlated with the microbial community (as above) or are of biological importance (TN, TOC, TON, and S were added for the latter). Letters indicate significant differences between groups based on an ANOVA test (p < 0.05); different letters indicate significant differences between those different sites. Units are as follows: temperature, ℃, conductivity, µS/cm; GWC (gravitational water content), TC (total carbon), TN (total nitrogen), TOC (total organic carbon), TON (total organic nitrogen), all %. All elemental values are given as ppb. TR: Tramway Ridge. WC: Western Crater.

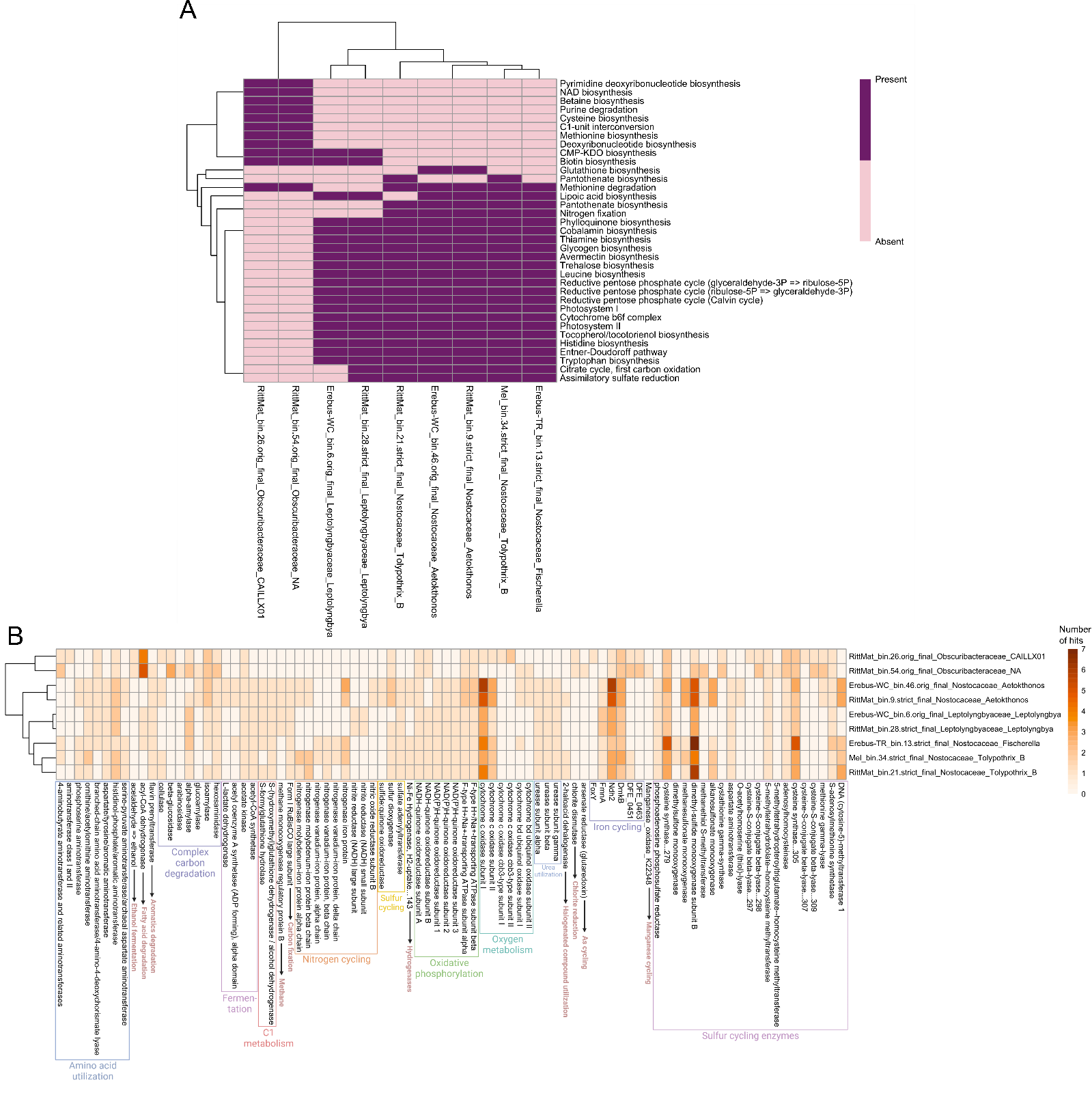

**Supplemental Figure 7** Functional comparison of Cyanobacterial MAGs as generated by METABOLIC showing (A) only the presence/absence of metabolic pathways and (B) the number of gene hits in the entire genomes. For (A), pathways that were absent from or present in all three Cyanobacteria groups were removed for ease of visualization.

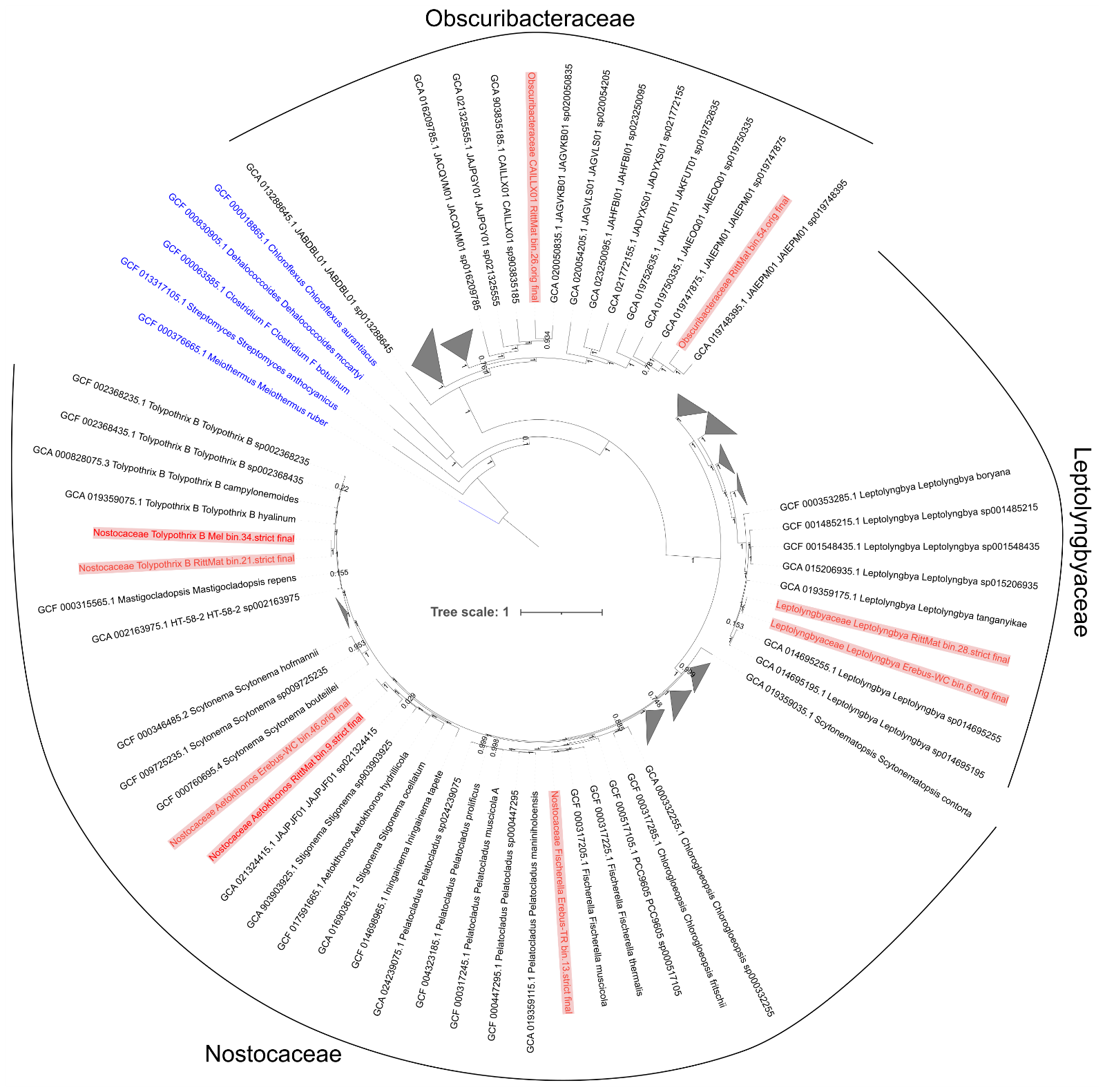

**Supplemental Figure 8** A rooted, approximately-maximum-likelihood phylogenomic tree of all cyanobacterial metagenome assembled genomes (MAGs) from this study (in red) with GTDB species cluster representative genome assemblies from the same three Cyanobacterial families. Numbers on each branch split refer to local support values calculated by FastTree. Five members of the Terrabacteria supergroup, highlighted in blue (*Streptomyces coelicolor*, *Clostridium botulinum*, *Chloroflexus* *aurantiacus*, *Dehalococcoides mccartyi*, and *Meiothermus ruber*) were used as the outgroups to root the tree. Bar, 1 substitution per nucleotide position.

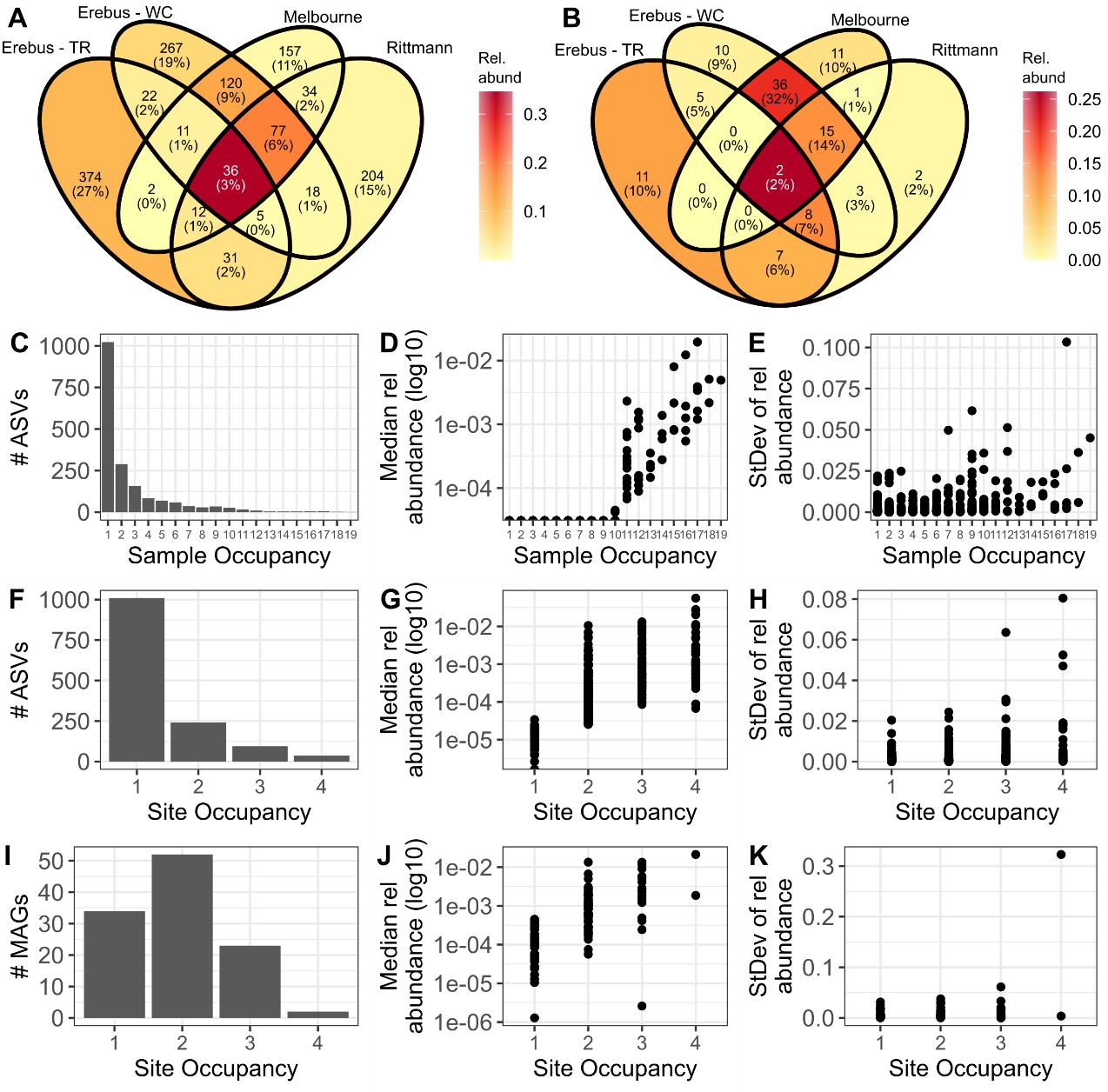

**Supplementary Figure 9** Exploring the common vs. site-specific microbes at these sites. (A, B) Venn diagrams comparing the number of (A) 16S rRNA gene sequence amplicon sequence variants (ASVs) or (B) metagenome assembled genomes (MAGs) unique or shared between different sites, colored by the combined relative abundance of species in each section. (C-K) The count (C, F, I), mean relative abundance (D, G, J), or standard deviation of relative abundance (E, H, K) of the number of (C-H) ASVs occupying different numbers of (C-E) samples or (F-H) sites, or (I-K) MAGs occupying different numbers of sites.

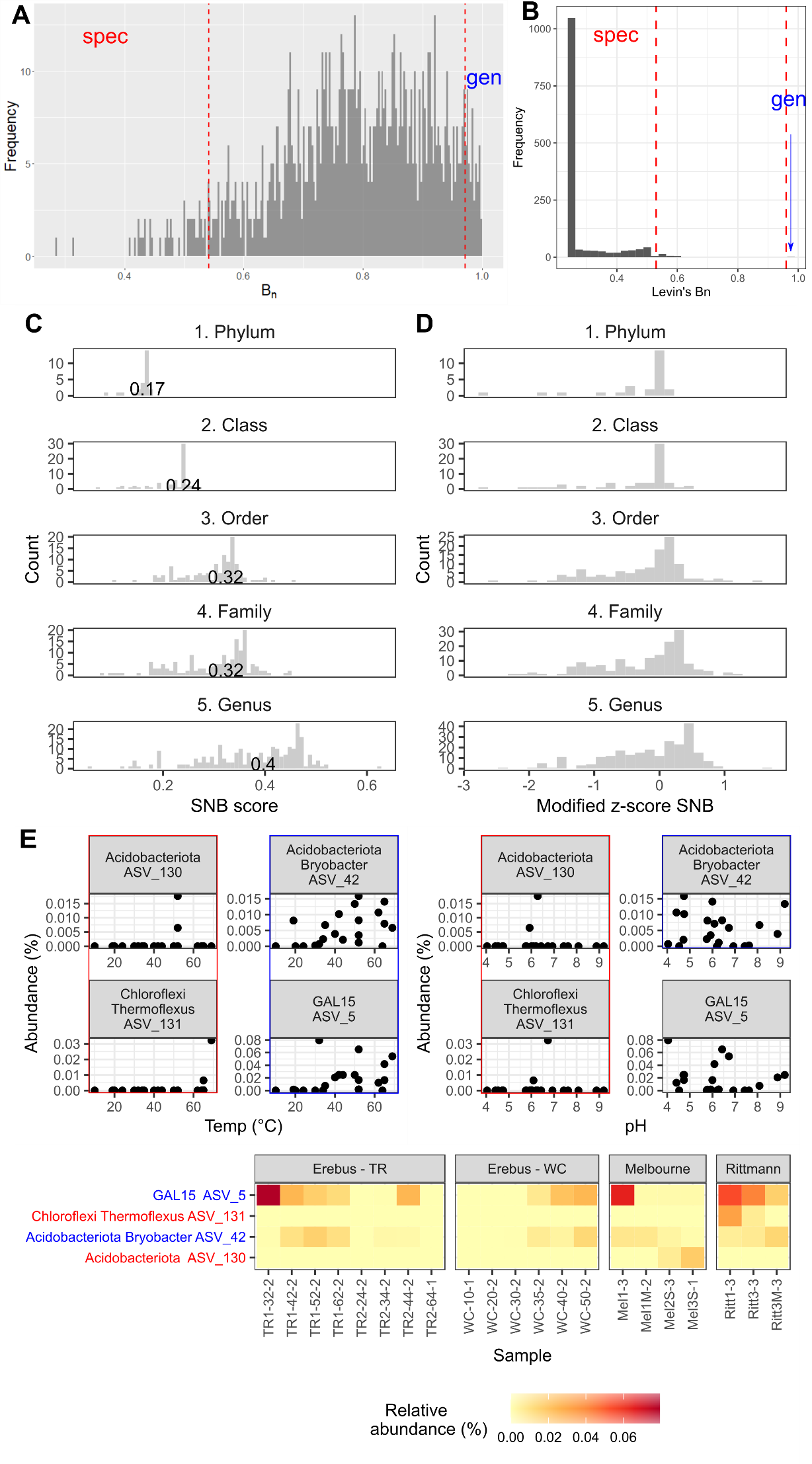

**Supplementary Figure 10** (A) Random theoretical distribution of Levin’s niche breadth (*B_N_*) produced as part of running the MicroNiche R package. (B) Frequency plots identifying the number of either specialists or generalists using *B_N_* on our 16S rRNA gene sequence data set, restricted to samples with a temperature above 35℃. (C-D) Social niche breadth (SNB) scores across different taxonomic levels in our data set. No Species level is given since the SNB calculations failed, given the large number of ASVs in our data set that were unassigned at the species level. (E) Plots showing the relationship between relative abundance and either temperature, pH, or site for either specialist or generalist ASVs based on variance across temperature and pH values, calculated using Hulbert’s niche breadth. ASV_5 did not qualify as a generalist for pH. The phylum and genus (if classifiable) are given for each ASV along with the ASV number. Specialist ASVs are highlighted in red, generalists in blue.

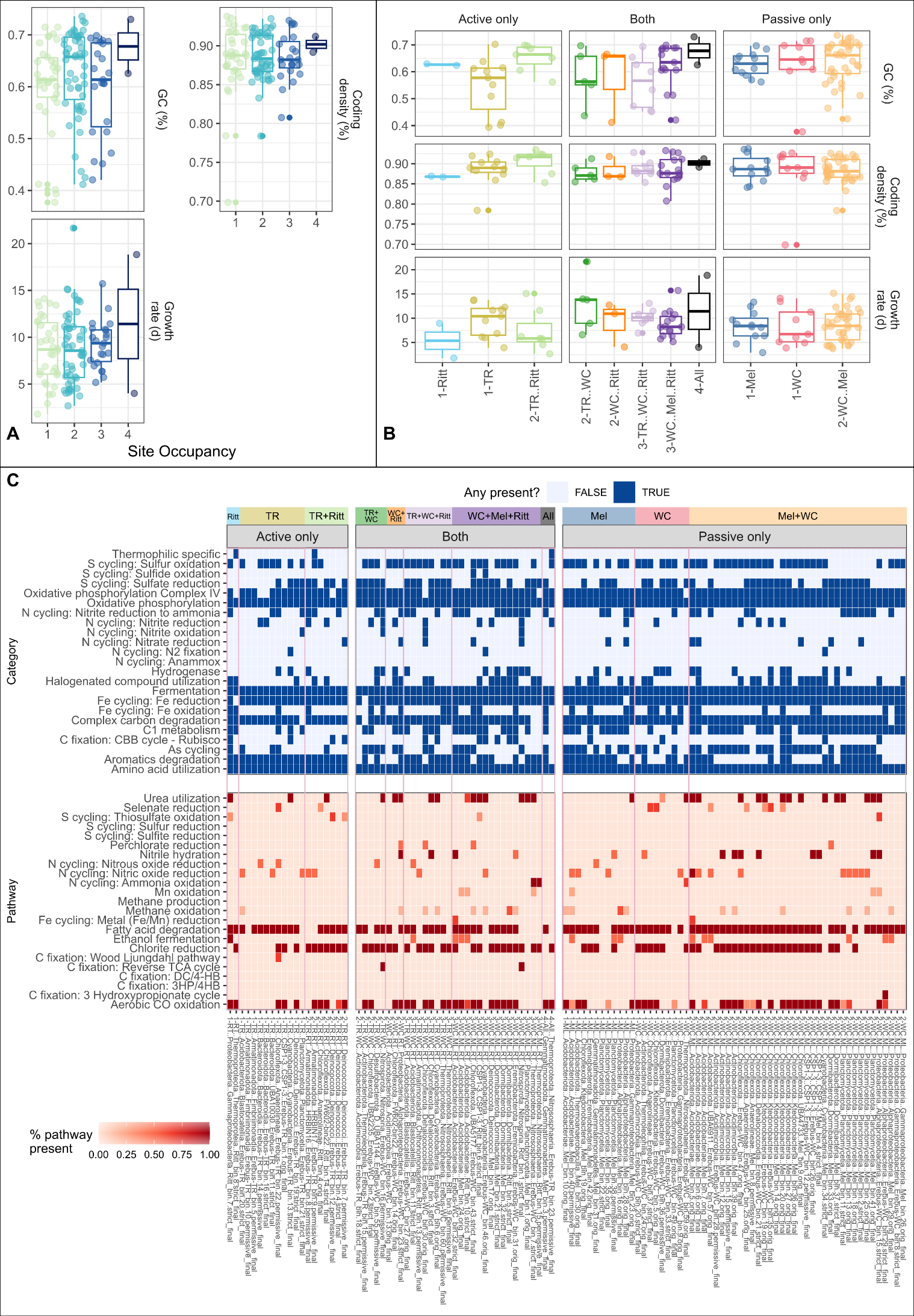

**Supplementary Figure 11** Comparison of genome characteristics between MAGs with different site occupancy patterns. No differences in any plots were significant (ANOVA, two-tailed). Ritt: Rittmann; TR: Erebus – Tramway Ridge; WC: Erebus – Western Crater; Mel: Melbourne. The number in front of the x-axis label in (B) represents the number of sites occupied by MAGs in that category. (C) Heatmap showing functional predictions for MAGs based on METABOLIC predictions, separated by site occupancy (restricted to one type of site, active or passive, or found in both types of sites) and whether the metabolic predictions are categories (containing multiple, non-sequential metabolic reactions) or pathways. Pink boxes differentiate different site occupancy patterns. X-axis MAGs are clustered first by site occupancy pattern, then by Phylum-Class. TR: Tramway Ridge. WC: Western Crater. RT: Rittmann. ML: Melbourne.
